# Co-translational folding of the first transmembrane domain of ABC-transporter CFTR is supported by assembly with the first cytosolic domain

**DOI:** 10.1101/2020.12.19.423590

**Authors:** Bertrand Kleizen, Marcel van Willigen, Marjolein Mijnders, Florence Peters, Magda Grudniewska, Tamara Hillenaar, Ann Thomas, Laurens Kooijman, Kathryn W. Peters, Raymond Frizzell, Peter van der Sluijs, Ineke Braakman

**Author notes:** These authors contributed equally (Marcel van Willigen, Marjolein Mijnders). Corresponding author: Ineke Braakman, PhD, Cellular Protein Chemistry, Faculty of Science, Utrecht University, Padualaan 8, 3584 CH Utrecht, The Netherlands, (with voice mail).

## Abstract

ABC-transporters transport a wealth of molecules across membranes and consist of transmembrane and cytosolic domains. Their activity cycle involves a tightly regulated and concerted domain choreography. Regulation is driven by the cytosolic domains and function by the transmembrane domains. Folding of these polytopic multidomain proteins to their functional state is a challenge for cells, which is mitigated by co-translational and sequential events. We here reveal the first stages of co-translational domain folding and assembly of CFTR, the ABC-transporter defective in the most abundant rare inherited disease cystic fibrosis. We have combined biosynthetic radiolabeling with protease-susceptibility assays and domain-specific antibodies. The most N-terminal domain, TMD1 (transmembrane domain 1), folds both its hydrophobic and soluble helices during translation: the transmembrane helices pack tightly and the cytosolic N- and C-termini assemble with the first cytosolic helical loop ICL1, leaving only ICL2 exposed. This N-C-ICL1 assembly is strengthened by two independent events: i) assembly of ICL1 with the N-terminal subdomain of the next domain, cytosolic NBD1 (nucleotide-binding domain 1); and ii) in the presence of corrector drug VX-809, which rescues cell-surface expression of a range of disease-causing CFTR mutants. Both lead to increased shielding of the CFTR N-terminus, and their additivity implies different modes of action. Early assembly of NBD1 and TMD1 is essential for CFTR folding and positions both domains for the required assembly with TMD2. Altogether, we have gained insights into this first, nucleating, VX-809-enhanced domain-assembly event during and immediately after CFTR translation, involving structures conserved in type-I ABC exporters.

## INTRODUCTION

Correct folding of proteins is essential for their biological function. Proteins fold largely co-translationally, from their N- to C-terminal ends. This helps to minimize the risk of aberrant intramolecular interactions as well as inappropriate encounters with simultaneously synthesized proteins on the polysome. The majority of eukaryotic proteins consist of multiple domains, yet most detailed folding studies were done on proteins consisting of single domains. As such, these studies provide limited insight into folding pathways of multidomain proteins, let alone membrane-spanning polytopic proteins. Folding starts co-translationally and individual domains of several multidomain membrane proteins have been shown to even complete their folding on the nascent chain [1, 2]. Principal questions include when and where in the folding pathway domains assemble; this can occur anywhere from nascent chain departure from the ribosomal exit tunnel to long after translation termination [3].

ABC transporters are amongst the most conserved and oldest protein superfamilies with members in all domains of life. They are multidomain multispanning membrane proteins that hydrolyze ATP to transport a variety of substrates (from nutrients, metabolites, vitamins, drugs, and lipids, to trace metals and small ions) across membranes. Their defects often cause disease [4]. All ABC transporters have a similar domain architecture with at least two transmembrane domains (TMDs) and two cytoplasmic nucleotide-binding domains (NBDs) [5, 6]. The ABC transporter CFTR forms a chloride channel across the apical plasma membrane of mainly respiratory and intestinal epithelial cells [7, 8]. Mutations in CFTR cause cystic fibrosis, the most common monogenic inherited and lethal disease [9, 10]. CFTR is a 1480-amino-acid polypeptide chain that constitutes an assembly of five domains (Figure 1A): two TMDs, each with six transmembrane helices that are connected through two intracellular helical loops (ICLs), and two NBDs. The fifth domain is not detected in the structure shown in Figure 1A; it is an intrinsically disordered regulatory region R, which undergoes phosphorylation cycles that regulate CFTR function. We showed before that the individual domains fold largely co-translationally in the ER [1]. Beautiful work from the Skach lab has shown that co-translational folding of NBD1 is optimized via tuning of local rates of translation and subdomain compaction [11]. The TMDs assemble via their TM and ICL helices and interact with the cytosolic NBDs through coupling helices at the ICL tips [12, 13]. Correct assembly of the individual domains is essential for CFTR channel function. This is underscored by analysis of F508del-CFTR, the most prevalent cystic fibrosis-causing mutation.

**Figure 1:**
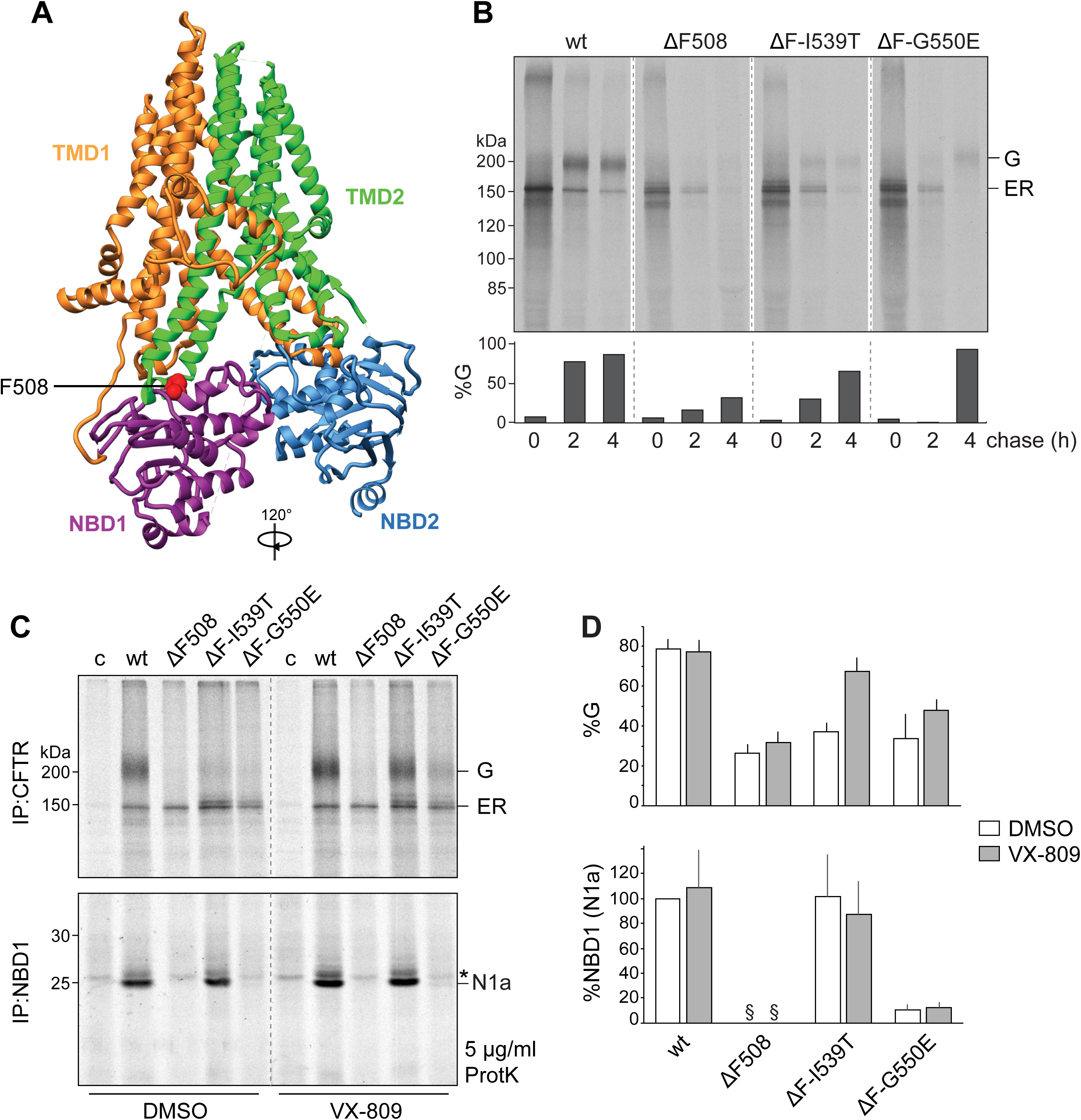
Intragenic suppressors enhance effect of VX-809 on CFTR biosynthesis. (A) CFTR structure image was made using the cryo-EM structure of CFTR (PDB: 5UAK) [12]. TMD1 in orange, NBD1 in purple, TMD2 in green, NBD2 in blue, R not shown. F508 in NBD1 in red. (B) HeLa cells expressing CFTR or the indicated CFTR mutants were pulse labeled for 15 minutes and chased for 0, 2, or 4 hours. CFTR was immunoprecipitated from 12.5% of the detergent lysate using MrPink antiserum against NBD1 and resolved on 8% SDS-PA gel. The graph shows quantifications of the Golgi-modified form (G) relative to total CFTR (ER + G) at each time point. (C) Similar as in (A) but chased for 2 hours only. When indicated, cells were pre-treated with VX-809 (3 μM) for 1 hour and kept present throughout the entire experiment, including lysis and proteolysis. CFTR was immunoprecipitated as in (B), and the remainder of the lysate was subjected to limited proteolysis after which protease-resistant NBD1 fragments were immunoprecipitated by anti-NBD1 (MrPink). Undigested CFTR was analyzed on 8% SDS-PA gel and the NBD1 protease-resistant fragments on 12% SDS-PA gel. Presence of the NBD1-specific fragment N1a indicates that NBD1 folded into a protease-resistant structure, whereas absence of fragment N1a indicates that NBD1 did not fold properly. (D) Quantifications of the Golgi-modified form (G) and folded NBD1 (N1a) relative to total CFTR (ER + G) from the same lane. Values of N1a are expressed as percentage of full-length protein, normalized to wild type in DMSO. wt, n=5; F508del (ΔF), n=5; F508del-I539T (ΔF-I539T), n=4; F508del-G550E (ΔF-G550E), n=3; error bars SEM. ER, ER form of CFTR; G, complex-glycosylated Golgi form of CFTR, which has passed the Golgi complex and may reside in or beyond the Golgi complex, including the plasma membrane; wt, wild-type CFTR; ΔF, F508del CFTR; c (control), transfection with empty vector; N1a, protease-resistant NBD1-specific fragment; *, background band; ^§^, below detection level.

The F508del mutation is located in NBD1 (Figure 1A), causes NBD1 misfolding and impairs expression at the plasma membrane [14–17]. F508del-induced NBD1 misfolding induces domain-assembly defects through disruption of the interaction of ICL4 in TMD2 with the F508 region in NBD1. As a result also the assembly of NBD2 with ICL2 in TMD1 is defective [18–20]. Second-site suppressor mutations in the F508del mutant protein may either restore NBD1 folding or rescue NBD1-TMD2 assembly, depending on their location within CFTR. Importantly, combinations of suppressor mutations that offset different defects in F508del CFTR have been shown to push folding and function of the mutant more effectively towards wild-type levels [21, 22].

Mechanistic insight in the folding of individual domains is limited. Even less is known of the timing and mechanism of domain assembly. In this study, we have analyzed early events in the folding and assembly pathway of CFTR under physiological conditions, using radiolabeling pulse-chase experiments in combination with limited proteolysis. We have focused on TMD1 and NBD1, because these are the two most N-terminal domains of CFTR, which are synthesized first and fold co-translationally [1]. Corrector compound VX-809 is a useful tool, because it was reported to stabilize TMD1 [23, 24] and enhance domain assembly, amongst others with NBD1 [25].

## RESULTS

To examine early events during biosynthesis of CFTR in the endoplasmic reticulum (ER), we radiolabeled live cells expressing CFTR. A short incubation with ^35^S-cysteine/methionine labeled a newly synthesized cohort of proteins (Figure 1B, 0-h chase). This was followed by a so-called chase, an incubation without radiolabel, to follow the labeled cohort over time. During the pulse and the 2-h and 4-h chase, the newly synthesized CFTR cohort folds and assembles its domains, and then is transported from the ER to the Golgi complex and cell surface. Upon arrival in the Golgi, the 2 N-linked glycans placed on CFTR during synthesis become modified, which is detectable as a decrease in electrophoretic mobility in SDS-PAGE (Figure 1B, wt, 2h and 4h of chase). As expected [17], ∼80% of wild-type CFTR had reached the Golgi complex in 4 h.

To assess the importance of NBD1, we also analyzed the F508del-CFTR mutant (ΔF508), which has a misfolded NBD1 domain [14, 15, 17, 26]. The fate of every CFTR variant is either degradation (when misfolded) or transport out of the ER to Golgi and cell surface. Of F508del CFTR ∼95% is degraded in 4 h [27, 28]; of the remaining 5%, ∼30% of the mutant had reached the Golgi complex (Figure 1B). We also used two intragenic suppressor mutations I539T or G550E [29–31], which improve folding of full-length F508del CFTR, slightly diminish degradation [17], and slightly increase transport to the Golgi (Figure 1B, increase in %G).

Folding of wild-type NBD1 and misfolding of F508del NBD1 is probed with a limited proteolytic digestion [1, 17]. More folded proteins are more resistant to digestion. Proteinase-K digestion of wild-type CFTR, followed by immunoprecipitation of NBD1 fragments, yields an ∼25-kDa protease-resistant NBD1 fragment. This fragment is protease sensitive and absent in F508del CFTR digests (Figure 1C, D) [17]. Only suppressor mutation I539T but not G550E completely restores folding of F508del NBD1 (Figure 1C, D) [17]. Despite restored folding of the F508del-I539T NBD1 domain, this was insufficient to effectively rescue the full-length protein from the ER (Figure 1B-D) [17].

### Intragenic suppressors enhance effect of corrector drug VX-809 on CFTR biosynthesis

CFTR folds its domains mostly co-translationally [1] and the F508del defect and rescue by I539T arise as soon as NBD1 is being synthesized [17]. We therefore reasoned that clinical corrector drug VX-809 (lumacaftor) [32, 33] may act early during CFTR biosynthesis and would serve as useful tool for our analysis of early domain folding and assembly. VX-809 did not increase protease resistance of NBD1, neither in any of the mutants nor in wild-type CFTR (Figure 1C, D). Exit of F508del CFTR from the ER was increased only slightly, as expected [32], but especially the I539T suppressor in F508del responded strongly to VX-809, with a rescue to ∼70% CFTR in the Golgi (Figure 1C, D). The synergy of VX-809 and I539T implied complementary rescue mechanisms: as I539T rescues folding of the NBD1 domain but not of full-length CFTR (Figure 1C, D, DMSO), we concluded that VX-809 must have salvaged domain assembly. In contrast to I539T, the primary folding defect in NBD1 is not rescued by G550E, nor by VX-809, and not by their combination either (Figure 1C, D). The strong synergy of G550E and VX-809 on full-length F508del CFTR with a misfolded NBD1 suggests that corrector drug and G550E must have improved domain assembly through two distinct domain interfaces. The folding of CFTR must involve at least three important phases: NBD1 folding (improved by I539T), and two domain-assembly interfaces, one improved by VX-809 (for which TMD1 is candidate) and one by G550E (for which NBD2 is candidate, considering the location of G550E in NBD1).

### VX-809 acts early during CFTR biosynthesis

To establish the timing of VX-809-susceptible domain assembly during CFTR folding, we used cells expressing F508del-I539T CFTR and added the drug at different times during the pulse-chase protocol (Figure 2A). As shown above, I539T alone provides minor escape from the ER (Figure 2A, lane 1), and this increased to 70% of wild-type levels with VX-809 present throughout the entire pulse-chase experiment. We set this maximum at 100% in Figure 2A, lanes 7 and 15). Addition of VX-809 at different stages of the pulse and chase (Figure 2A) showed that later addition led to a gradual decrease in rescue. With VX-809 present during pulse labeling only, transport from the ER already was at ∼80% of maximal rescue (Figure 2A, lanes 9 and 16), showing that the compound acted predominantly during and early after translation. Pre-treatment before pulse labeling was without effect on F508del-I539T (Figure 2A, lane 11, and compare lanes 12 and 16), showing that VX-809 was washed out effectively during the 15-min starve incubation, and that the effect on CFTR was not due to alterations of the cellular proteome.

**Figure 2:**
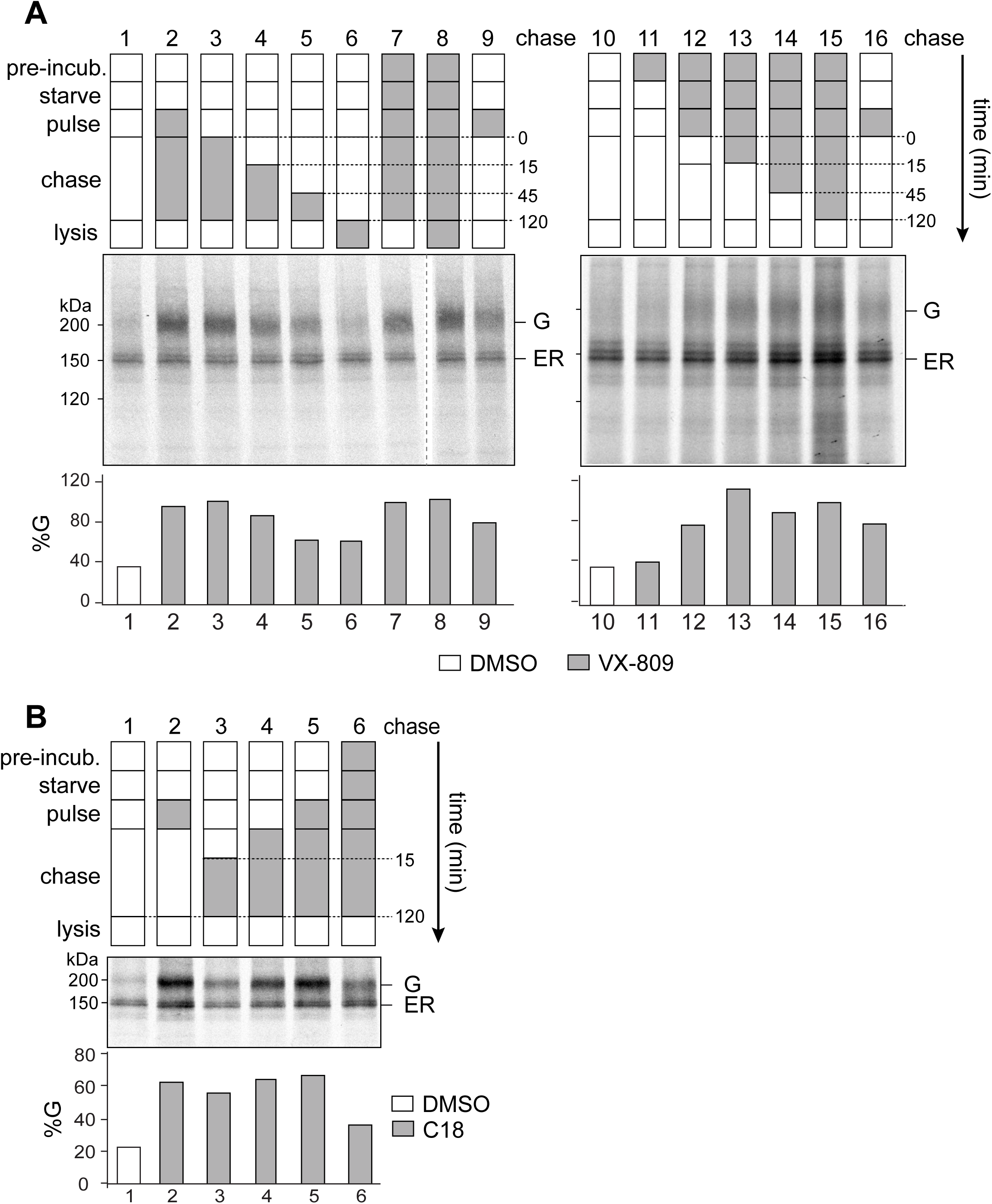
VX-809 acts early during CFTR biosynthesis. (A) HeLa cells expressing F508del-I539T-CFTR were pulse labeled for 15 minutes, chased for 2 hours, and lysed. VX-809 (3 μM) was present in the indicated phases (grey boxes). When VX-809 was included in the lysis buffer, it was kept present during limited proteolysis as well. Lanes 1-9 and 10-16 are from two separate, representative experiments, in which the VX-809-induced rescue was related to the maximal rescue in lanes 7 and 15, set at 100%. The graph shows quantifications of the Golgi-modified form (G) relative to total CFTR (ER + G) in the same lane. ER, ER form; G, complex-glycosylated Golgi form, which has passed the Golgi complex and may reside in or beyond Golgi, including the plasma membrane.

The broad heterogeneity in both translation and folding rates within the CFTR population did not allow a more precise determination of the time window of VX-809 activity on F508del-I539T-CFTR. These heterogeneities lead to nascent CFTR chains being finished far into the chase, long after incorporation of radiolabel has stopped [34]. This explains the significant correction seen when VX-809 was present only during the chase. Similar results were found using VX-809 analog C18 (Figure 2B). We conclude that VX-809 and C18 improved domain assembly at an early folding phase without restoration of NBD1 misfolding. The effect may be co- and/or post-translational, possibly coincident with the co-translational folding of NBD1, misfolding of F508del, and restoration by I539T [17]. Because VX-809 does not act on cells before CFTR is synthesized, we confirmed that the corrector likely targets CFTR itself rather than another cellular protein, as suggested before [23, 24, 32].

### VX-809 increases protease resistance of TMD1, but not of NBD1 and TMD2

To identify the conformational basis for the improved exit of CFTR from the ER, we examined the effect of VX-809 on CFTR domains. We in-vitro translated CFTR and parts of CFTR: TMD1 (E395X), TMD2 (837-1202X), as well as CFTR truncated after NBD1 (D674X) or after R (E838X) (Figure 3A). For explanation of construct nomenclature see Materials & Methods. In E383X, residue 383 is replaced by a stop, leading to a construct with C-terminal residue 382 (Figure 3A). The in-vitro synthesis was done in the presence of ^35^S-labeled amino acids and ER membranes derived from digitonin-permeabilized cells, and in the presence or absence of corrector drug (Figure 3B, C). Limited proteolysis to probe CFTR conformation here yields all protease-resistant fragments from the translated CFTR construct, without the bias of immunoprecipitation [1, 17]..

**Figure 3.**
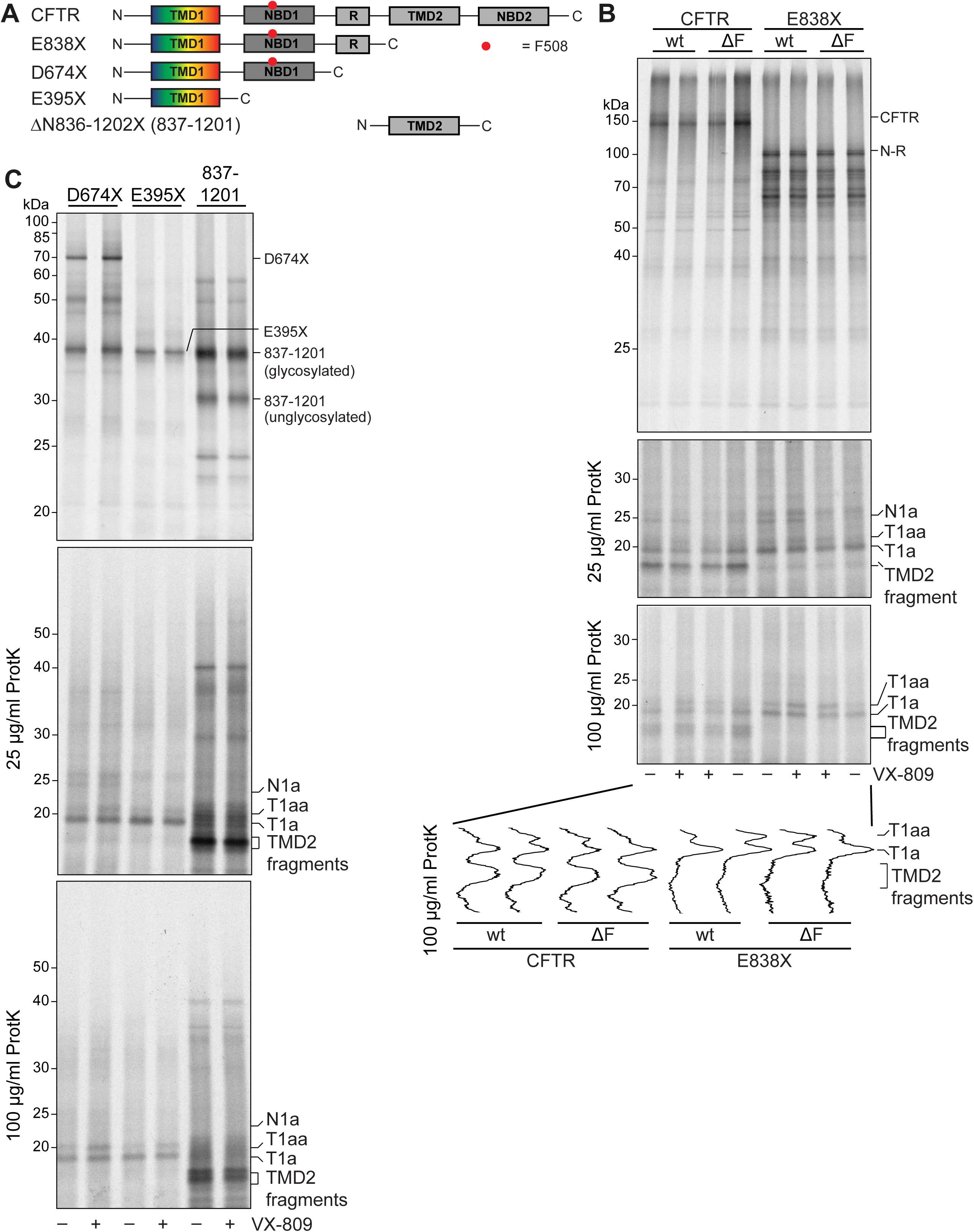
VX-809 acts on TMD1 but not NBD1 and TMD2 from in-vitro-translated CFTR. (A) Schematic representation of the constructs used in this figure. For explanation of construct nomenclature see Materials & Methods. (B) CFTR or CFTR truncated downstream of the R-region (E838X), wild-type or F508del, were translated and translocated in vitro in the presence of semi-intact cells as source of ER membranes for one hour at 30 °C with or without 5 μM VX-809 and analyzed by 12% SDS-PAGE (top panel). The bottom panels show protease-resistant fragments after digestion with two concentrations of Proteinase K analyzed by 12% SDS-PAGE. The lane-intensity profiles (ImageQuant analysis) of protease-resistant fragments T1a and T1aa of the 100 μg/mL Proteinase-K digests are shown below. (C) Similar experiment as in (B), for shorter constructs as indicated. Top panel shows undigested translation products and bottom panels show the Proteinase-K digests. wt, wild-type CFTR; ΔF, F508del CFTR; T1a and T1aa indicate protease-resistant TMD1 fragments; N1a indicates protease-resistant NBD1 fragment.

The corrector drug did not have an effect on the quantity of protein translated. Digestion of CFTR with 25 μg/mL Proteinase K yields relatively protease-resistant fragments for NBD1 (N1a), TMD1 (T1a and T1aa), and TMD2 [1] (Figure 3B, C). Removing TMD2 and NBD2 from the translated construct (E838X) led to loss of the TMD2 fragments and an increase in detectable NBD1 and TMD1 fragments (Figure 3B), perhaps because more of the shorter protein was produced. F508del in both proteins leads to loss of the N1a fragment [1] (Figure 3B). Upon increasing protease concentration to 100 μg/mL, a clear VX-809-related change appeared in the proteolytic profiles of CFTR and E838X, independent of F508del background: an increase in T1aa (Figure 3B). At this protease concentration, most NBD and R fragments are degraded while TMD fragments remain [1]. NBD1 and TMD2 fragments did not show a response to VX-809 (Figure 3B).

To limit the number of proteolytic fragments, we translated isolated TMD1 (E395X), TMD2 (837-1202X), and a TMD1-NBD1 construct (D674X) (Figures 3C and S1A). The proteolytic profiles (in particular at 100 μg/mL Proteinase K) demonstrated that isolated TMD1 (E395X in Figure 3C) displayed the same VX-809-related structural change as TMD1 in full-length and the other truncated forms of CFTR, E838X in Figure 3B, and D674X in Figure 3C: an increase in the T1aa fragment whereas T1a was the same with and without VX-809. Again, TMD2 was not affected by VX-809.

In summary, we showed that NBD1 (N1a) and TMD2 (Figure 3B, C) did not respond to VX-809, and found a change only in TMD1 fragments (Figure 3B, T1a and T1aa).

### Conformational rescue of F508del arises from protection of the N-terminus of TMD1

Through unbiased in-vitro experiments we now had established that TMD1 was the primary target of VX-809 (Figure 3), consistent with previous reports [23, 24, 35]. To confirm the biochemical effect of VX-809 on TMD1 in intact cells, we pulse labeled cells expressing TMD1 (E395X; Figure 4A) for 20 min in the absence or presence of VX-809 and/or Corrector 4a (C4) [36] (Figure 4B). C4 was added as negative control for the effect of VX-809 on TMD1, because it stabilizes TMD2 [35, 37] and hence acts additively with VX-809 on F508del-CFTR [38]. Protease-resistant fragments were immunoprecipitated using an antibody against ECL1 [39] (Figure 4A). Neither VX-809 nor C4 changed the levels of radiolabeled TMD1/E395X (Figure 4B, top panel, lanes 10-12). As seen in vitro (Figure 3B, C), T1aa increased upon VX-809 but not upon C4 treatment of cells (Figure 4B, bottom gel panel and bar graph of the ratio T1aa/T1a), identical to the VX-809-effected change in in-vitro translated TMD1/E395X (Figure 3C).

**Figure 4:**
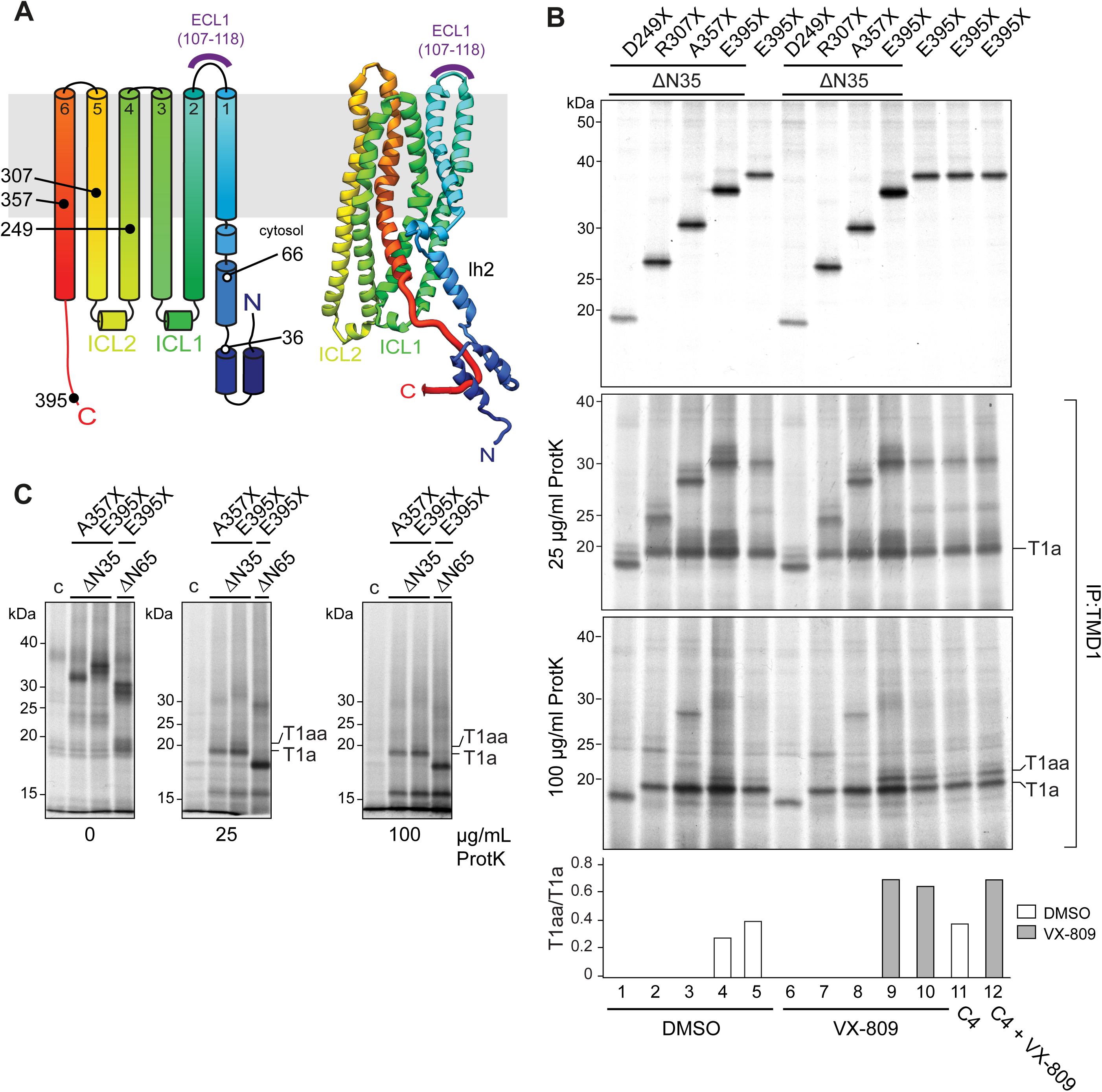
Protease resistance of TMD1 characterized. (A) Schematic and structural rainbow representation of TMD1 indicating the location of the ECL1-antibody epitope (purple) and the boundaries of N- and C-terminally truncated constructs used here. We used a CFTR model [42], as it is more likely to represent the structure of the TMD1 N-terminus during translation. (B) CFTR truncations, with boundaries indicated in (A), were expressed in HEK293T cells and radiolabeled for 20 min. Where indicated, 10 μM corrector VX-809 and/or 10 μM Corrector 4a (C4) was present before and during the labeling. The top panel shows translated proteins immunoprecipitated using the E1-22 antibody against ECL1 in TMD1 and analyzed by 12% SDS-PAGE. The bottom panels show TMD1 fragments immunoprecipitated by the ECL1 antibody after digestion with 25 or 100 μg/mL Proteinase K, analyzed by 12% SDS-PAGE. The graph shows the T1aa/T1a ratios of the 100 μg/mL Proteinase-K digests. Constructs either have or lack the N-terminal 35 residues. To avoid confusion, we annotated in the Figure the N-terminal and C-terminal mutations separately: ΔN35-D249X then is 36-248, ΔN35-R307X is 36-306, ΔN35-A357X is 36-356, and ΔN35-E395X is 36-394, In the Results text we used the names of either the N- or the C-terminal truncations, again to avoid confusion as to which end of the construct or fragment is being described. (C) CFTR truncations were in-vitro translated and translocated in the presence of semi-intact cells as source of ER membranes for one hour at 30 °C and analyzed by 12% SDS-PAGE. The two right panels show protease-resistant fragments after proteolysis with Proteinase K analyzed by 12% SDS-PAGE. Nomenclature as in panel (B): ΔN35-A357X then is 36-356, ΔN35-E395X is 36-394, and ΔN65-E395X is 66-394. c (control), transfection with empty vector; T1a and T1aa indicate protease-resistant TMD1 fragments. For explanation of construct nomenclature see Materials & Methods.

TMD1 consistently yields an ∼19-kDa fragment we called T1a (Figure 4B; [26]). The stabilizing effect of VX-809 on TMD1 always led to appearance or increase of T1aa, slightly larger than T1a (Figures 3B, C, 4B bar graph), and sometimes to an increase in T1a as well, together showing increased protease resistance of TMD1. To identify T1a and T1aa, mass spectrometry is no option because it lacks the requisite sensitivity to detect the trace amounts of radiolabeled fragments in the samples. We therefore analyzed T1a and T1aa from cells expressing a range of N- and C-terminally truncated TMD1 constructs.

Truncating 65 but not 35 amino acids (Figures 4B, C) from the N-terminus led to a shift down of T1a, implying that the N-terminus of the T1a was between 35 and 65. Zooming in on this region (Figure 5A) then showed that removing 50 residues was too much (it led to a T1a-like fragment that ran as too small) whereas removing 49 residues precisely generated T1a (Figure 5B). Similarly, C-terminal truncations showed that replacing E257 for a stop codon generated T1a (Figure 5C) whereas all shorter constructs led to a downward shift of T1a (Figures 4B, 5C). T1a thus represents TMD1 fragment 50-256, with Leu 49 and Ser 256 as the most prominent cleavage sites (Figure 6A).

**Figure 5.**
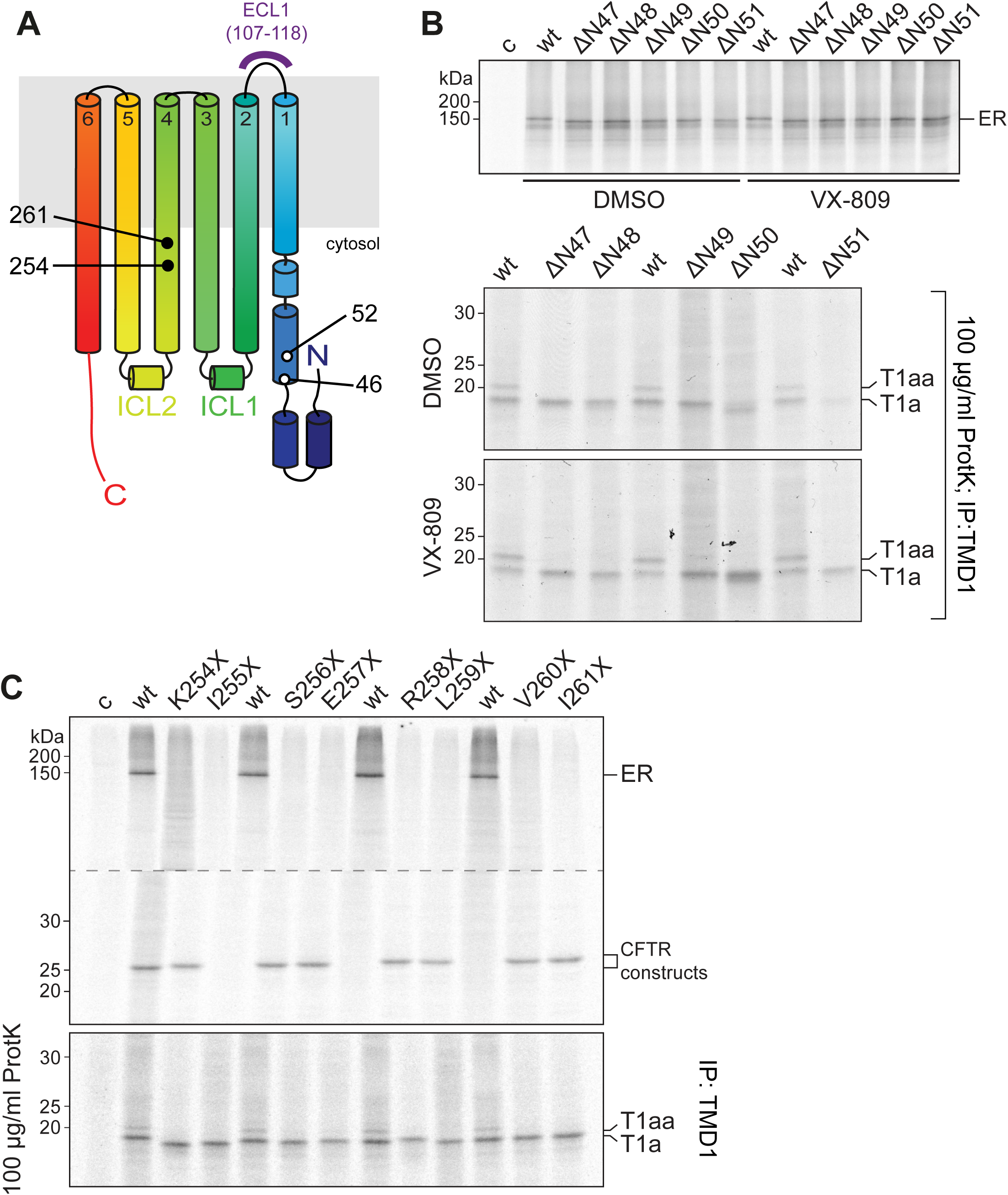
Protease-resistant TMD1 fragment T1a identified. (A) Schematic rainbow representation of TMD1 indicating the location of the ECL1 antibody epitope (purple) and the boundaries of N- and C-terminally truncated constructs used here. We used a CFTR model [42], as it is more likely to represent the structure of the TMD1 N-terminus during translation. (B) CFTR truncations, with bounding residues indicated in (A), were expressed in HEK293T cells and radiolabeled for 20 min, with or without corrector VX-809 (10 μM) present before and during the labeling. The top panel shows undigested protein immunoprecipitated with MrPink. The bottom panels show TMD1 fragments immunoprecipitated with the E1-22 antibody against ECL1 after digestion with 100 μg/mL Proteinase K. (C) As in (B), without VX-809. The top panel shows undigested protein immunoprecipitated with MrPink. The bottom panel shows TMD1 fragments immunoprecipitated with the ECL1 antibody after digestion with 100 μg/mL Proteinase K. (C)-(D) show that T1a represents TMD1 fragment 50-256. ER, ER form; T1a and T1aa indicate protease-resistant TMD1 fragments; wt, wild-type CFTR; c (control), transfection with empty vector.

**Figure 6.**
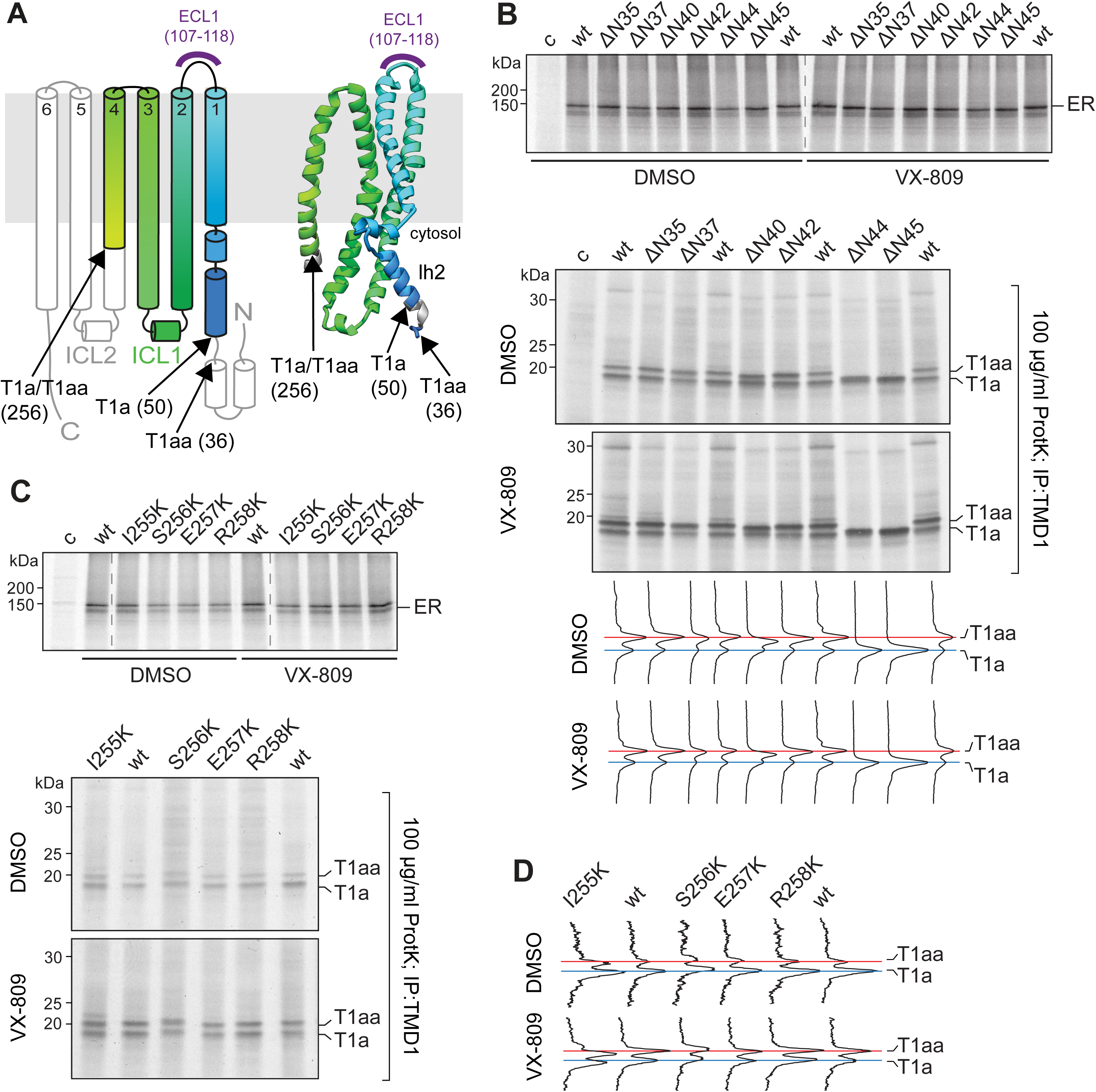
Conformational rescue of F508del arises from protection of the N-terminus of TMD1. (A) Identity of the T1a and T1aa fragments depicted in cartoon and structure representation as in Figure 4A. (B) HEK293T cells expressing indicated constructs were pulse labeled for 20 min. Where indicated, corrector VX-809 (10 μM) was present before and during the labeling. The top panels show undigested protein immunoprecipitated with MrPink. The bottom panels show TMD1 fragments immunoprecipitated with E1-22 after digestion with 100 μg/mL Proteinase K. The lane profiles are of the 100 μg/mL Proteinase-K digests. T1aa/T1a quantifications are shown in Figure S2. (C) Similar experiment as in (B), for shorter constructs as indicated. Top panel shows undigested translation products and bottom panels show the Proteinase-K digests. (B)-(C) show that T1aa represents TMD1 fragment 36-256 (see panel (A)). (D) Lane profiles of the 100 μg/mL Proteinase K digestion gel of panel (C). T1aa/T1a quantifications are shown in Figure S2. ER, ER form; T1a and T1aa indicate protease-resistant TMD1 fragments; wt, wild-type CFTR; c, (control), transfection with empty vector. Blue and red lines show electrophoretic mobilities of wild-type T1a and T1aa, respectively. Data are representative of 3 independent observations.

As for T1a, we established that the most likely N-terminus of T1aa is residue 36 (Figure 6A, B). Removing 37 N-terminal residues but not 35 caused a downward mobility shift of T1aa (Figure 6B): the red line marks the position of wild-type T1aa; it helps visualize differences in peak positions. Further truncations led to a larger shift, with T1aa from ΔN44 and ΔN45 already collapsed onto T1a. T1aa most likely resulted from Proteinase-K cleavage after Ser 35 and not Asp 36 because the protease does not cleave behind Asp [40]. Deviating from the series of increasing downward mobility shifts was ΔN42: ΔN37 shows the first shifted fragment, ΔN40 yields an again smaller fragment, and ΔN44 and larger truncations do not yield a detectable T1aa anymore as its mobility has merged with T1a. The odd behavior of ΔN42 may be due to the loss of proline 41, as prolines are known to influence mobility: its loss may have changed local conformation and SDS binding. Any alternative cleavage site does not allow a plausible scenario that is consistent with all data. Ser 42 for instance is a possible Proteinase-K cleavage site but this would leave the shift down of T1aa from constructs ΔN37 and ΔN40 unexplained and would imply that the full shift from T1aa to T1a would be realized between ΔN42 and ΔN44. Further truncations (Figure 6B) do not yield any T1aa either. Wild-type T1a is marked with a blue line.

To establish the C-terminus of T1aa we used a different strategy, as T1aa was lost already when TMD1 was truncated C-terminally from E395X down to A357X (Figure 4B). The C-terminal boundary of T1aa cannot be between 357 and 395, because this would imply that T1aa would be at least 357-256=101 residues larger than T1a (when extended at the C-terminus alone). Because the N-terminal extension of 14 residues is sufficient to explain the difference between T1a and T1aa, we examined whether these fragments shared their C-termini and whether the loss of T1aa from A357X would require an alternative explanation. Electrophoretic mobility often changes when a charged residue is added or removed. We therefore used a limited lysine scan around the predicted C-terminus of T1a, mutating 1 residue at a time into a lysine. Again a blue line was used to mark wild-type T1a and a red line for wild-type T1aa. I255K and S256K both yielded a shift in T1a as well as in T1aa compared to wild-type fragments (Figure 6C, D). The I255K-derived fragments ran slightly lower, the S256K-derived fragments slightly higher than wild type, whereas mutating residues 257 and higher did not lead to a shift (Figure 6C, D). This confirmed that T1a and T1aa had the same C-terminus, and that 256 but not 257 and 258 were part of both fragments (Figure 6C, D). T1a hence constitutes residues 50-256, and T1aa most likely 36-256. As the signature of the VX-809 effect on CFTR was an increase in T1aa, we concluded that VX-809 improved CFTR conformation by effecting an increased protection/shielding of Leu 49 at the N-terminal edge of Lasso helix (Lh) 2 in the N-terminus of TMD1. The T1aa/T1a ratio was increased in every VX-809-responsive construct (bar graphs in Figures 3, 4, 6, and S2).

### The N- and C-termini of TMD1 are essential for cytoplasmic-loop packing in CFTR folding

With the identification of T1a and T1aa the question remained why the fragments from 357X did not include T1aa. The A357X construct lacks the entire C-terminal TMD1-NBD1 linker region (Figure 7A). Refining the truncation series showed that C-terminal residues 376-385 in TMD1 were essential for generation of T1aa (Figure 7B), for protection of Leucine 49 in the N-terminus. As neither T1a (50-256) nor T1aa (36-256) contained these residues, the C-terminus of TMD1 must have exerted its influence at a distance. The CFTR structures [12, 13, 41] and models [42, 43] indeed show that residues 376-385 tightly pack with N-terminal helices (Figure 7C, blue and red), and that the N- and C-termini of TMD1 tightly associate with the first intracellular loop (ICL1) (Figure 7C, green). We therefore set out to determine whether ICL1 was needed for this N-C packing and for the improvement by VX-809 (Figure 7C), and designed mutations aimed to disrupt the interface. We added, reverted, or removed charges in K166 to V171 in ICL1, which is close to residues 376-385 in the TMD1 C-terminus (Figure 7C). Mutations K166Q and R170G were found in CF-patients (http://www.genet.sickkids.on.ca) but were not confirmed to be CF-causing (https://cftr2.org) [44].

**Figure 7.**
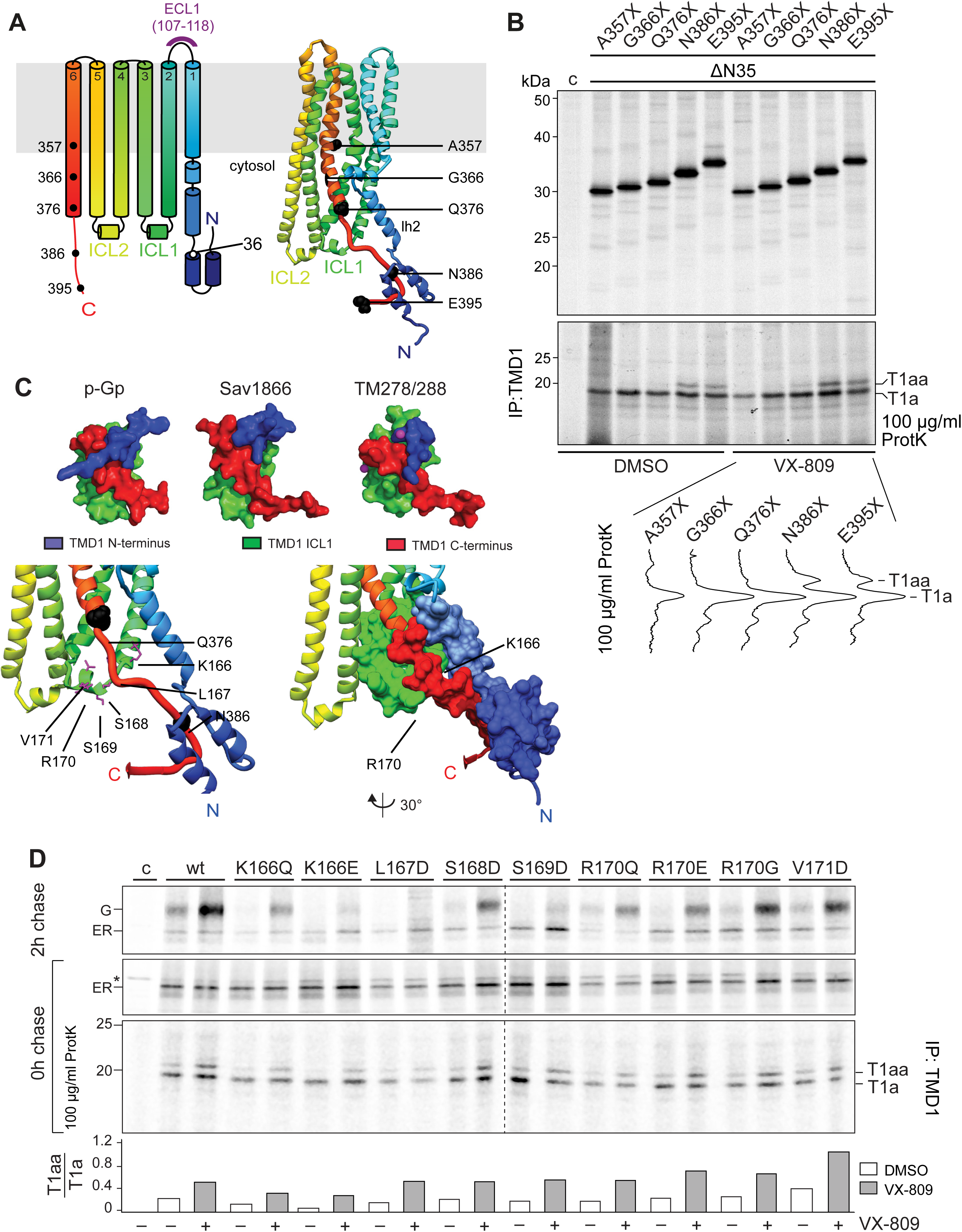
Evolutionarily conserved packing of TMD1 is important for CFTR folding and transport to the cell surface. (A) Schematic and structural rainbow representation of the N-terminus and TMD1 indicating the location of the ECL1 (also E1-22) antibody epitope (purple) and the boundaries of C-terminally truncated constructs used here. (B) HEK293T cells expressing indicated constructs were pulse labeled for 20 min. Where indicated, corrector VX-809 (10 μM) was present before and during the labeling. The top panel shows undigested protein immunoprecipitated with anti-TMD1 antibody ECL1. The bottom panel shows TMD1 fragments immunoprecipitated with ECL1 after digestion with 100 μg/mL Proteinase K. T1a and T1aa indicate protease-resistant TMD1-specific fragments. Lane profiles show the relative intensities of the two fragments. T1aa/T1a quantifications are shown in Figure S2. All constructs lack the N-terminal 35 residues. To avoid confusion, we annotated in the Figure the N-terminal and C-terminal mutations separately: ΔN35-A357X then is 36-356, ΔN35-G366X is 36-365, ΔN35-Q376X is 36-375, ΔN35-N386X is 36-385, and ΔN35-E395X is 36-394, In the Results text we used the names of either the N- or the C-terminal truncations, again to avoid confusion as to which end of the construct or fragment is being described. (C) Top, structure of ICL1 (green), the N- (blue) and C-termini (red) in surface representation of TMD1 in different type-I ABC exporter classes. Human CFTR, PDB: 5UAK [12]; P-glycoprotein, mouse (M.musculus), PDB: 4Q9H [83]; Sav1866, S. aureus, PDB: 2HYD [59]; TM278/288, T. maritima, PDB: 3QF4 [84]. Bottom, structures representing the packing of residue stretch 376-385 in TMD1’s C-terminus (red) with the ICL1 (green) and the N-terminus (blue), on the left in ribbon representation, on the right partly as surface representation. The residues in ICL1, from K166 to V171, were mutated and analyzed by pulse chase and limited proteolysis in (D). (D) HEK293T cells expressing CFTR without or with mutations in ICL1 were pulse labeled for 15 minutes and chased for 2 hours. When indicated, 3 μM corrector VX-809 was present before and during the pulse and chase. The top two panels show undigested samples immunoprecipitated using antibody 2.3-5 against NBD2; the bottom panel shows protease-resistant fragments immunoprecipitated with the E1-22 against ECL1 antibody after digestion with 100 μg/mL Proteinase K. The bar graph shows the T1aa/T1a ratio of the 100 μg/mL Proteinase-K digests. ER, ER form; G, complex-glycosylated Golgi form, which has passed the Golgi complex and may reside in or beyond Golgi, including the plasma membrane; T1a and T1aa indicate protease-resistant TMD1 fragments; wt, wild-type CFTR; *, background band; c (control), transfection with empty vector.

We expressed these mutants in HEK293T cells, pulse labeled for 20 min, chased for zero or two hours, both in the absence or presence of VX-809, and subjected the detergent lysates to limited proteolysis. All mutants showed a defect in transport to the Golgi complex: mutants K166Q, K166E, L167D, and S169D, R170E were most defective, whereas S168D, R170G, and V171D showed a milder defect (Figures 7D, 8A). This interface thus must be important for exit from the ER. Fragment T1a appeared in all mutants, indicating that early folding of TMD1 was not affected.

**Figure 8.**
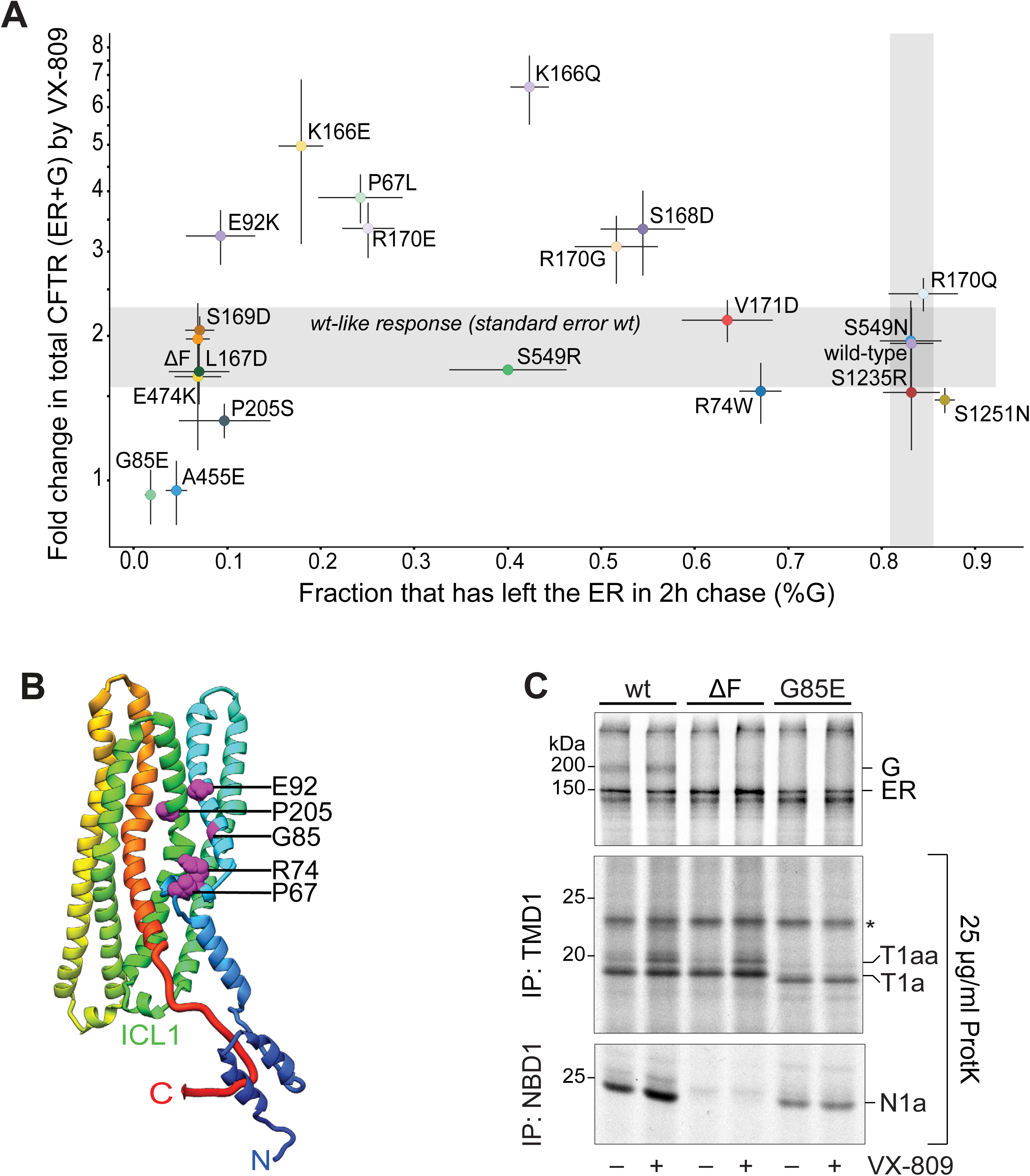
Comparison identifies CFTR mutants hyper- and hypo-responding to VX-809. (A) Fold change in (mutant) CFTR expression (ER+G) by VX-809 (y-axis) plotted against the fraction that had left the ER during 2 hours of chase (G/(ER+G) in the DMSO control samples (x-axis). Quantifications were obtained from experiments as in Figure 7D; n≥3 for each mutation. The horizontal and vertical grey zones show the standard error for wild type: ‘wt-like response’. The cut off for the ‘wt-like VX-809 response’ is 1.5x SEM of wt-CFTR samples (n=8). Using the ‘wt-like’ cut off we separated high-VX-809 responders from low-VX-809 responders. (B) Structural representation (Cryo-EM, PDB: 5UAK) in rainbow of TMD1, highlighting TMD1 residues analyzed in (A), all found mutated in CF patients except E92. (C) HEK293T cells expressing CFTR or two CF-causing CFTR mutants were pulse labeled for 15 minutes and chased for 2 hours. When indicated, 3 µM corrector VX-809 was present before and during the pulse and chase. The top panel shows undigested samples immunoprecipitated using NBD2-specific antibody 2.3-5. The lower panels show TMD1- and NBD1-specific protease-resistant fragments immunoprecipitated with the ECL1 antibody and MrPink respectively, after digestion with 25 µg/mL Proteinase K. T1aa/T1a quantifications are shown in Figure S2. (B) and (C) show that T1aa represents TMD1 fragment 36-256. ER, ER form; G, complex-glycosylated Golgi form, which has passed the Golgi complex and may reside in or beyond Golgi, including the plasma membrane; T1a and T1aa indicate protease-resistant TMD1 fragments; wt, wild-type CFTR; *, background band.

All constructs tested in Figures 7B and D responded to VX-809 with an increase in ratio T1aa/T1a (Figures 7D and S2). To compare VX-809 responses, the fold increase in transport to the Golgi was calculated from the sum of ER and Golgi because the fraction transported to the Golgi often was too low in control conditions to generate reliable ratios. As the post-ER fate of CFTR mutants is either transport to the Golgi complex or degradation, the sum of ER and Golgi forms are not only a proxy for transport, but also for stability. Each mutant responded to VX-809, often similar to wild-type CFTR, with an ∼2-fold increase in total protein (Figure 7D, 8A). Striking was the hyper-response of the S168D, R170G, and R170E mutants (∼3-fold), and especially the K166 mutants (∼5-6-fold more CFTR). This hyper-response also led to a strong increase In T1aa from the K166 mutants. Whereas this fragment is usually visible after pulse labeling, it barely appeared in these mutants, which shows that Leu 49 was less protected in these mutants. The hyper-response suggested that the K166 mutants represented the folding defect that was restored best by VX-809.

We have shown that ICL1, and especially K166, is crucial for the tight packing of TMD1, and that VX-809 causes strongest rescue of mutants defective in this packing. For comparison we added CF-disease-causing mutants to the analysis (The Clinical and Functional TRanslation of CFTR (CFTR2); available at https://cftr2.org) [44], and some that initially but erroneously were suspected to be disease causing (http://www.genet.sickkids.on.ca, underlined), in TMD1 (P67L, R74W, G85E, E92K, <u>L177T</u>, P205S), NBD1 (A455E, E474K, F508del, S549N), or NBD2 (<u>S1235R</u>, S1251N). Especially P67L, R74W, G85E, E92K, L177T, and P205S are likely important for the helical packing of TMD1 (Figure 8B, magenta spheres). Limited digestion with Proteinase K showed that all tested mutants yielded the typical T1a fragment, indicative of native-like TMD1 folding, except G85E, whose ‘T1a’ fragment had shifted mobility. G85E also was the exception in not yielding a T1aa fragment upon VX-809 treatment (Figure 8C), whereas all other mutants tested showed increased T1aa quantity (Figures 8C and S2). G85E does not respond to VX-809 most likely because of its defective insertion into the ER membrane [45, 46]. We concluded that all tested mutants except G85E responded to VX-809 by increased protection of the N-terminus.

The improved packing of TMD1 by VX-809 did not always lead to enhanced transport to the Golgi (Figure 7D, 8A), implying that the improved folding was not the bottleneck for export of these mutants. Many mutants were in the wild-type range of response (Figure 8A, grey zones in x- and y-axes), even the more severe folding mutations such as F508del-CFTR. Two mutant groups deviated: the low/non-responders and the hyper-responders. Strong responders –the K166 mutations, P67L, and E92K–all located in the area where TMD1 N- and C-termini pack with ICL1 (Figure 8B). E92K in TM1 was shown before to respond well to VX-809 [24], perhaps because it affects the TMD1 packing VX-809 rescues.

When transport to the Golgi cannot be rescued by VX-809 (Figure 8A) three scenarios may be at play: i) the mutation may prevent drug binding; ii) VX-809 may bind but may not improve TMD1 packing due to the defect; or iii) VX-809 works but the CFTR defect requires different or additional help (such as perhaps P205S and A455E). ICL1 mutant L167D did not respond much in terms of exit from the ER, although T1aa did increase upon VX-809 treatment (Figure 7D). Patient mutant E474K is located on the NBD1 surface and may disrupt interaction with ICL1, amongst others with R170 [12, 13, 41], which may explain its responsiveness.

We concluded that packing of the N- and C-termini onto ICL1 is important for early TMD1 folding in CFTR, which is likely conserved in other ABC exporters, such as mouse (M.musculus) P-glycoprotein, S. aureus Sav1866, and T. maritima TM278/288, shown in Figure 7C. Corrector VX-809 improves TMD1 packing, acts on TMD1 alone and thus must bind to TMD1 as well.

### Effects of NBD1 and corrector drug on TMD1 are additive

Immediately after synthesis of TMD1, NBD1 is translated and may natively interact with ICL1. To examine whether NBD1 contributes to the folding/packing of TMD1, we compared proteolytic stability of TMD1 alone (1-394) with that of TMD1-NBD1 (1-673) expressed in HEK293T cells and analyzed as above. Upon addition of NBD1 to the construct, the T1aa fragment increased, an effect similar to that of VX-809 (Figure 9B). Combining NBD1 and VX-809 increased T1aa further (Figure 9B), indicating an additive effect of NBD1 and VX-809 on TMD1, especially visible in the bar graph of T1aa/T1a ratio), an effect we confirmed in in-vitro translation experiments (Figures 3C, S1A, B, bar graph in S1B). The additive effect suggested a distinct mechanism with similar outcome: increased protection of the N-terminus of TMD1. VX-809 analogue C18 displayed the same effects (Figure S1C, again especially visible in the bar graph).

**Figure 9.**
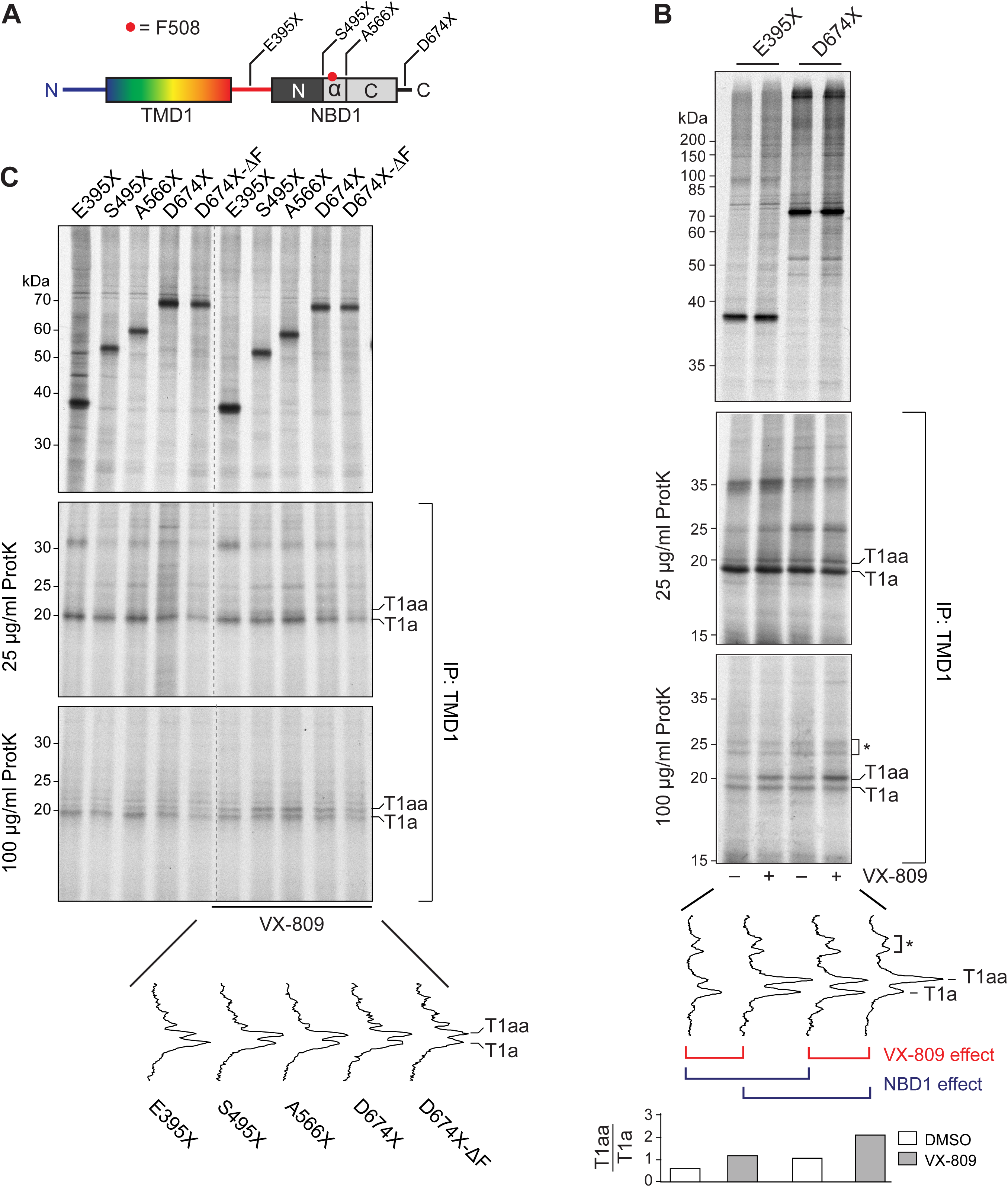
N-terminal subdomain of NBD1 and corrector improve on TMD1 folding additively. (A) A schematic representation of C-terminal truncations; TMD1 in rainbow and the three NBD1 subdomains from dark to light grey. (B) HEK293T cells expressing indicated truncations were pulse labeled for 20 min in the presence or absence of 10 μM VX-809. The undigested samples in the left panel were immunoprecipitated by ECL1. The right panels show TMD1-specific protease-resistant fragments immunoprecipitated by the ECL1 antibody after digestion with 25 or 100 μg/mL Proteinase K. The lane profiles of the 100 μg/mL Proteinase-K digests are shown. The graph shows the T1aa/T1a ratio of the 100 μg/mL Proteinase-K digests. See also Figure S1B for in-vitro confirmation. (C) HEK293T cells expressing indicated truncations were pulse labeled for 20 min in the presence of 10 μM VX-809 control. When indicated, the radiolabeled lysates were subjected to 25, or 100 μg/ml Proteinase K, after which TMD1-resistant fragments were immunoprecipitated using the ECL1 antibody. All samples were analyzed by 12% SDS-PAGE. Lane profiles of the 100 μg/mL Proteinase-K digests are shown. T1aa/T1a quantifications of these are in Figure S2. ΔF, F508del CFTR; T1a and T1aa indicate protease-resistant TMD1 fragments; *, background bands.

### N-terminal subdomain of NBD1, residues 389-494, are sufficient for TMD1 folding

The effect of NBD1 on the N-terminus of TMD1 was suggestive of early assembly of TMD1 with NBD1. If so, perhaps only a part of NBD1 would be sufficient to exert this effect. We therefore set out to identify the minimal part of NBD1 that would affect TMD1 folding. We examined two additional constructs truncated within NBD1 (resembling nascent chains released from the ribosome): one just before the α-helical subdomain (S495X), which contains only the N-terminal (α–β) subdomain (TMD1 plus 100 residues from NBD1) and one immediately after the α-helical subdomain (A566X) (Figure 9A), the region in NBD1 that not only contains the F508 residue, but also suppressor mutation I539T, which rescues F508del-NBD1 folding. Lane intensity profiles of 100 µg/ml Proteinase-K digests (Figure 9C) and the T1aa/T1a bar graph (Figure S2) clearly showed that already the shortest C-terminal NBD1 truncation (S495X) displayed the NBD1-dependent folding effect on TMD1. The F508 deletion (Figures 9C and S2), which is located in the α-helical subdomain of NBD1, slightly enhanced this effect, which confirmed that only the N-terminal subdomain of NBD1 was sufficient to increase protease protection of the N-terminus of TMD1, to improve co-translational folding of TMD1.

## DISCUSSION

The folding of polytopic multispanning membrane proteins such as ABC-transporters is a major challenge. We here present the physiological solution to this conundrum: the start of folding is co-translational and modular. Most of the individual-domain folding occurs co-translationally [1, 47], which is facilitated by the relatively slow translation of CFTR [48]. Packing of the most N-terminal domain, TMD1, occurs independent of other domains. It is enhanced by early assembly of the downstream polypeptide stretch, the N-terminal subdomain of the cytosolic NBD1 domain. That corrector drug VX-809 improves TMD1 packing as well, through a different mechanism and additive with NBD1, supports the conclusion that this first domain-assembly event forms an essential early phase in the folding of such a complex protein.

Early folding of individual domains in multidomain proteins such as CFTR starts co-translationally [1, 47]. We have provided experimental evidence that TMD1 and NBD1 interact already on released nascent chains, which implies interaction during synthesis. The limited-proteolysis read out for early TMD1 folding (T1a) was found in the isolated domain as well as the full-length protein, in both in-vitro translations and in intact cell experiments. The T1a protease-resistant fragment contains TM1-TM4 packed with the N-terminal Lasso helix 2 (cleaved at residue L49, starting at residue 50), and a small part of ICL2 (to residue 256). Tight packing of N- and C-termini of TMD1 led to increased protection of Leu 49 in Lh2, which was enhanced by corrector drug VX-809 and by addition of NBD1, in additive fashion with different mechanisms (Figure 10, steps 2-4). This TMD1-NBD1-linker-dependent Leu 49 protection led to extension of the protease-resistant TMD1 fragment with 14 amino acids upstream of Lh2 (T1aa); this revealed increased protease resistance of the TMD1 N-terminus (Figure 10, events 3-4). Enhanced packing of TMD1 required only the N-terminal part of NBD1, and not the F508-containing α-helical subdomain; it therefore was independent of proper NBD1 folding (Figure 10, events 3-4). This first domain-assembly step of TMD1 with NBD1 positions NBD1 for interaction with ICL4 in TMD2, likely stimulating TMD1-TMD2 assembly (Figure 10, event 5). Of note is that most CF-causing mutations are located in the N-terminal half of CFTR, TMD1 and NBD1 [44]. As this structure is highly conserved in type I ABC-exporters, the CFTR folding pathway may be general for this class.

**Figure 10.**
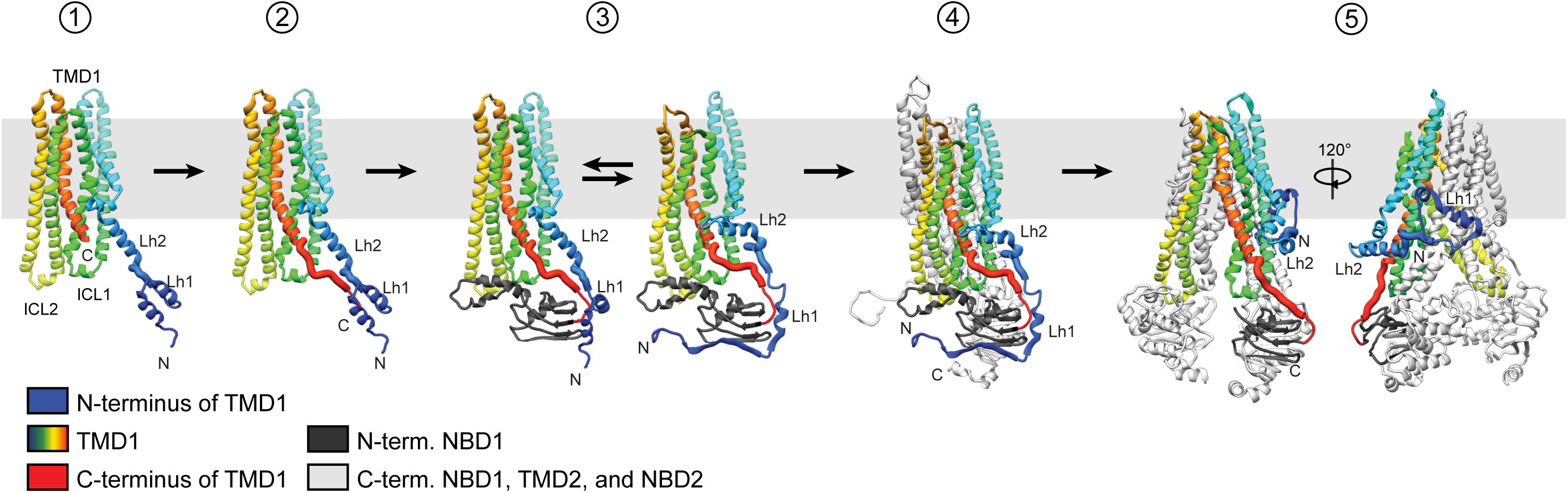
Proposed model for co-translational packing of the N- and C-termini and ICL1 of TMD1, with NBD1. Schematic model of the proposed TMD1 folding mechanism. TMD1 folds largely co-translationally. Structural images 1-4 were made using models of CFTR from Callebaut and colleagues with the CFTR N-terminus in the cytoplasm [43]. Synthesis of the C-terminus of TMD1 (red) induces its packing with ICL1 (green) and the N-terminal lasso helix 2 (Lh2; blue) (from 1 to 2). This packing is improved by expression of the N-terminal subdomain of NBD1 (dark grey) and by VX-809 (3). The packed TMD1 interface provides a template for further CFTR folding and domain assembly, hence explaining the VX-809 mode of action on TMD1, facilitating downstream folding events (4). The N-terminal lasso structure containing lasso helix 1 (Lh1) may form after TMD2 synthesis (4-5). Image 5 was made using the cryo-EM structure of CFTR (PDB: 5UAK) [12], in which the N-terminus is partly embedded in the bilayer and interacts with TMD2. Image 5 is represented in Figure 1A with different color coding for the individual domains.

### TMD1 folding: TMD1 packing

Other studies have focused on TMD1 packing and assembly with other domains, which were performed using cross-linking techniques in steady state [23, 25]. Here we show the kinetics of this TMD1 folding process using radiolabeling of newly synthesized CFTR, with different folding events represented by changed protease-resistance of the N-terminal half of TMD1. The protease-resistant TMD1 fragment suggests that a typical trimeric helical bundle is formed between TM1-3, a basic and evolutionarily conserved folding unit found in multispanning membrane proteins [49]. In the closed-channel cryo-EM structure [13], TM4 has more distance from the trimeric TM1-3 core. This would explain the Proteinase-K accessibility of serine 256 in ICL2 and suggests that early TMD1 packing may well resemble a native-like structure. This is confirmed by the co-translational formation of T1a [1].

Increased co-translational packing of TMD1, promoted by the corrector drug VX-809 and the N-terminal part of NBD1, increased protease resistance of Lasso helix 2. The VX-809 effect was dependent on the C-terminus of TMD1, the linker between TMD1 and NBD1, which indicates increased tightness of packing of N- and C-termini with ICL1. The protection induced by the N-terminus of NBD1 may be the result of an interaction between ICL1 and NBD1, which would stabilize TMD1 folding, as suggested by various CFTR structures [12, 13, 41] and models [42, 43]. Alternatively, the long TMD1 N-terminus may be stabilized by interaction with the N-terminus of NBD1 [43]. Recently published cryo-EM structures show the CFTR N-terminus along the membrane, embracing TMD1 and TMD2 [12, 13] (step 5, Figure 10). Before synthesis of TMD2 and assembly of TMD1 and TMD2, a location of the CFTR N-terminus in the cytosol associating with NBD1 is not unlikely. This was shown in various structural models [42, 43]. The N-terminal lasso motif of CFTR may well adopt multiple conformations during folding, or even during its functional cycle, moving in and out of cytosol and membrane, interacting with different parts of CFTR, including regulatory region R [50].

In-vivo studies in *E. coli* demonstrated for several membrane proteins that specific ‘packing’ interactions between multiple N-terminal TM segments and a C-terminal TM segment already occurred during translation, as soon as the C-terminal helix was synthesized; these are key to a stable tertiary structure during translation [51]. TMD1 folding appears to be a similar co-translational event in which the N-terminal part awaits residues 376-385 located in the C-terminal TMD1-NBD1 linker to gain stability, with the difference that for TMD1 much of the packing occurs in the cytosol rather than within the bilayer. To allow this interaction, TM6 must ‘wedge’ between the TM1-3 trimeric TM bundle and TM4, which explains the protease sensitivity of ICL2 close to the membrane and increasing protease resistance of the TMD1 N-terminus. Cryo-EM structures suggest that TMD2 packing is conserved [12, 52, 53] and TMD2 therefore may fold and pack in a similar fashion as TMD1.

The lateral packing of TM helices in TMD1 is structurally conserved in all ABC-type I exporters and often may be guided by electrostatic interactions. K166 is only one example of a highly conserved, charged residue in ABC-transporters. Charge reversal of the K166 residue into glutamic acid in the coupling helix of ICL1 is causing a severe CFTR folding defect, which is rescued very well by VX-809. In all CFTR structures, both cryo-EM and 3D models, this K166 residue is sandwiched between the N-terminal Lh2 and the C-terminal TMD1-NBD1 linker region (residues 376-385). Two hydrophobic residues, F374 and L375 were reported to be crucial for stability of isolated TMD1 in cells [24]. We found that the mere presence of those residues was not sufficient for TMD1 folding, nor for the VX-809-induced TMD1 packing (Figure 6, see Q376X).

Joining of N- and C-terminal stretches is a common occurrence in folded domains and proteins [3, 51, 54], and must be important for stability of TMD1 (and likely TMD2) during the co-translational folding of CFTR. To propagate ion conductance, TM segments need charged residues embedded in the bilayer at the expense of their hydrophobicity and hence their individual stability in the membrane. TM6 is crucial for CFTR’s ion conductance [55], but on its own is unstable in the ER membrane [56]. Similarly, TM1 contributes to ion conductance [57], is unstable in the bilayer [58] and needs neighboring TM2 to co-translationally integrate in the bilayer during synthesis [45]. The ‘joining’ of N- and C-terminal elements, as we found for TMD1, facilitates to keep unstable TM segments in register during synthesis, *en route* to the native structure. For ABC transporters such as CFTR, TMD1 needs to maintain a ‘folding-competent’ state until TMD2 is synthesized and integrated into the membrane, since some ‘swapping’ of TM segments and their intracellular loops between both TMDs is needed for its native structure and for function.

### TMD1 folding: joining with NBD1 during translation

The first ABC-transporter crystal structures showed that the long helical intracellular loops (ICLs) from the TMDs anchor the NBDs to relay ATP hydrolysis allosterically to pore opening and closing [59, 60]. We here showed the importance of the ICL1-NBD1 interaction during folding. Only the N-terminal α–β subdomain of NBD1, which needs ATP for folding during translation [61], was required to improve TMD1 packing and folding (Figure 10, steps 3-4). As a consequence, co-translational, ‘vertical’ packing of the N-terminal part of NBD1 to the cytoplasmic parts of TMD1 assures TM6 stability [56], forms a template for the consecutive co-translational folding of NBD1 [11, 17], and promotes folding of the following CFTR domains.

The conclusion that the folding events we have uncovered occur during translation has the following basis: both TMD1 folding and NBD1 folding (and misfolding and rescue) occur on the nascent chain [1], and the biochemical effects of VX-809 and the N-terminus of NBD1 occur early and on the shortest constructs that mimick released nascent chains. Upon synthesis of the minimal 495 residues that show both effects (S495X, Figure 9A), CFTR translation has another ∼1,000 residues to go in intact cells, which at the slow rate of 2.7 aa/s [48] will take on average 6 min. It is extremely unlikely that the TMD1-NBD1 interfaces will hold off interaction until chain termination.

Several structural [12, 41, 42] and biochemical studies on CFTR confirmed that ICL1 from TMD1 connects to NBD1 [23, 25, 62, 63], and ICL2 from TMD1 ‘crosses’ to connect with NBD2 [62, 64], which explains its protease sensitivity. NMR studies showed that the ICL1-NBD1 interaction is dynamic and phosphorylation dependent [65], implying a role for ICL1-NBD1 in channel activity. We found that ICL1 residues 166-171 are at the center of domain interactions: laterally within TMD1, vertically with NBD1, together packing the N-terminal half of CFTR. Our biochemical data is consistent with the recent cryo-EM structures showing that the part of ICL1 we disrupted interacts with part of the N-terminal lasso motif, the TMD1-NBD1 linker region, and with elements in the N-terminal region of NBD1 [12]. A synthetic ICL1 peptide upstream of K166 was found to have highest binding affinity to purified NBD1 [63], suggesting that the TMD1-stabilizing effect of the NBD1 α–β subdomain may be caused mostly by interactions of this subdomain with the N-terminus of TMD1 rather than ICL1. Linear peptides lacking essential secondary structure may not fully represent the in-vivo situation, or the available structures represent only one of many conformations (and domain interfaces) CFTR may have.

Our conclusion that the lateral packing of N- and C-termini of TMD1 with ICL1 and the vertical packing of the NBD1 α–β subdomain with ICL1 and the TMD1 N-terminus are important for CFTR folding is based on several observations: i) Many mutations in these interfaces severely impact ER export of CFTR; ii) Mutants responding strongly to VX-809 are in the cytoplasmic packing interface of TMD1: P67L (this manuscript) [66] and L69H [67] in the N-terminal lasso motif, K166E/Q, S168D, and R170E/G in ICL1 (this manuscript), and the F374A and L375A in the TMD1-NBD1 linker region [24]; iii) Structural analysis shows conservation of these lateral and vertical interfaces in most other ABC-transporters.

Despite the importance of these TMD1-NBD1 interfaces for CFTR folding, stability, and VX-809-dependent correction, reversing or removing single charges in this area hardly affected proteolytic susceptibility that reads out for co-translational TMD1 folding; the T1a fragment was always present, and T1aa mostly as well. We concluded that TMD1 always adopts a similar core conformation that is not grossly disturbed by mutations. This is in line with our previous work showing that the CFTR domains fold mostly co-translationally and independently [1]. Structures and structural models such as in Figures 1A and 10 confirm that the interaction interfaces of TMD1 and NBD1 show touching of domains instead of entangling or strand swapping. Data presented here show that even the most severe mutations located in either TMD1 (G85E) or NBD1 (F508del) did not affect the other domain. This also explains why VX-809, which improves co-translational TMD1 folding, does not have any impact on NBD1 folding and stability as shown by our protease-susceptibility data and work by others [38]. That VX-809 promotes cross-linking of TMD1 to NBD1 [25] confirms our conclusion that the drug strengthens TMD1-NBD1 domain assembly through improvement of TMD1 folding rather than NBD1 domain folding.

### How VX-809 corrects F508del-CFTR

The F508del-CFTR mutation suffers from multiple molecular defects that change the cellular fate of the protein [68]: i) structural differences in F508del mRNA due to the codon change at position 507 [69, 70]; ii) co-translational intra-domain folding defect in NBD1 [17]; iii) impaired interdomain assembly, especially between NBD1 and TMD2 through ICL4 [21, 22]; iv) increased protein instability [14, 15] and turnover at the cell surface [71, 72].

Both the G550E suppressor [17, 73] and VX-809 show that the co-translational folding defect in F508del NBD1 does not need to be rescued to restore F508del-CFTR trafficking to the cell surface, implying that improved or restored domain assembly is the alternative to NBD1 rescue. Improved co-translational and early post-translational TMD1-NBD1 packing indeed leads to exit of F508del-CFTR from the ER. We here showed that VX-809 improves co-translational TMD1 folding, which improves TMD1 stability [25], and then strengthens the interaction of TMD1 with NBD1 [25]. VX-809 also promotes cysteine cross-linking between TMD1 (R170C) and NBD1-R-TMD2-NBD2 constructs expressed ‘in trans’ [25]. We found R170E and G mutations to be hyper-responders to VX-809. Interestingly, recent modeling efforts by Callebaut and co-workers suggested that residues ∼13-34, just downstream of Lh2, ‘wraps’ against the α–β-subdomain of NBD1 [42, 43]. This likely explains the enhanced early protease protection of Leu 49 in Lh2.

The domain assembly of NBD1 with TMD2 through ICL4 is described to be the most important interaction to improve F508del-CFTR folding and stability [21, 22]. Both the cryo-EM structures and the 3D models show that residues S168D and S169D in the coupling helix of ICL1 are within 4Å distance with the ‘378-QEY-380’ stretch from the C-terminal TMD1-NBD1 linker region, but also with the E474 acidic residue in the N-terminal α–β subdomain of NBD1. The E474 residue is also within 4Å range of W1063 and R1066 residues in the coupling helix of ICL4, adjacent to and even facing ICL1. Bioinformatic analysis of ABC-B and ABC-C transporters showed that residue E474 co-evolved with R1066, stressing the importance of this interaction [74]. The final step to local stability then is the connection between the ICL4 R1070 region and the F508 region in NBD1 [21, 22]. TMD1 interacting through both its N-terminus and ICL1 with the α–β subdomain of NBD1 likely occurs during translation. NBD1 then folds, assembling its subdomains, which positions NBD1 for interaction with ICL4. F508del NBD1 fails to fold and position the 507-509 region properly to ICL4, probably leading to failed assembly of TMD1 and TMD2. This positioning of NBD1 for ICL4 binding is in line with the observation that ICL4 binds much weaker than ICL1 to purified NBD1 ‘in trans’ [63].

This chain of events explains how VX-809-induced early packing of TMD1 leads to formation of a template for ICL4 to bind to, leading to progression of domain assembly in the absence of folded F508del-NBD1. This allostery in the VX-809 effects, as recognized before [75], also explains the previously suggested stabilization of the NBD1-ICL4 interface [24, 35, 38, 75, 76], the NBD1-ICL1 interface [25] and the TMD1-TMD2 interaction [35]. All may well be the consequence of improved TMD1 packing effected by VX-809. This extensive allostery causes the site of VX-809 binding to deviate from the many sites in CFTR where VX-809 shows impact.

### How, where and when does VX-809 act: site-of-binding vs. site-of-action

Despite the many changes induced by VX-809 (improvement of various domain interfaces and improved export from the ER), our studies showed that TMD1 was the only domain affected in isolation and the most likely direct recipient of the VX-809 effect. Strongly responding mutants, such as P67L, K166E/Q, S168D, and R170E/G, all reside in TMD1, which implies that VX-809 fully restores the effects of these mutants, suggesting that the binding site is close to this region. In-silico predictions suggested that the K166 residue is key in binding VX-809 or its analogs VX-661 and C18 [77]. If so, the binding site must contain other residues as well, because our K166 mutants (to E or Q) showed a strong response rather than the lack of response expected upon disruption of the drug binding site. To determine the VX-809 binding site, we may need to await high-resolution structures of CFTR complexed with the drug (reviewed in [78]).

Our data show that the lack of ER-to-Golgi trafficking cannot be equated to lack of a response to the drug. The two-fold rescue of exit from the ER implies that when a mutant responds, this response will only be detectable when the original (so-called residual) activity of the mutant is high enough to detect the change: 2x 1% of wild-type activity =2% and still too low to detect, but 2×10% = 20% and detectable. All mutants we have examined except for a few respond biochemically in TMD1 to VX-809, with an increase of T1aa. This would suggest that VX-809 is an effective component of a corrector drug cocktail for most CF-causing mutants. An exception is non-responding mutant G85E, which has defective TM1 insertion and stability [46], precluding rescue by improved packing of these misaligned TM helices.

Our radiolabeling approaches on nascent chain-mimicking constructs show that VX-809 acts during or soon after CFTR synthesis. Studies on purified CFTR reconstituted in proteoliposomes however demonstrated a C18-dependent effect on CFTR function [79]. VX-809 may bind to any form of CFTR that is not tightly packed, whether in purified form or transiently in the cell. CFTR needs the Hsp70/90 chaperone machines for first-time folding but also for stability at the cell surface, suggesting that its channel-opening-and-closing cycle may destabilize CFTR or that a metastable state is crucial for CFTR’s function [80]. This makes it likely that corrector VX-809, while acting predominantly early, can impact protein stability in vitro and at the cell surface as well.

## MATERIALS & METHODS

### Cell culture

HeLa cells (ATCC) were maintained in MEM supplemented with 10% fetal bovine serum (FBS), South America origin, 100 µM Non-Essential Amino Acids and 2 mM Glutamax. HT1080 fibrosarcoma cells were grown in DMEM supplemented with 8% FBS and 2 mM Glutamax, and HEK293T cells (ATCC) were maintained in DMEM containing 10% FBS and 2 mM Glutamax. Cells were cultured in humidified incubators at 37 °C in 5% CO_2_. All reagents were purchased from Life technologies. Figure 1 shows data from HeLa cells, whereas the other figures with live cells report on data with HEK293 cells. In-vitro translations use HT1080 or HEK293 cells as source of ER membrane. We use these human cell lines interchangeably to ensure generality of findings and preclude cell-line-specific effects.

### Antibodies and reagents

Generation of the polyclonal rabbit antiserum MrPink against purified hNBD1 was described before [17], as was polyclonal rabbit antibody ECL1, directed against the cyclic peptide of the extracellular loop 1 [39]. In figures 4B, 5B, 6B, C, and 7B its successor was used, polyclonal antibody E1-22, directed against the same epitope, designed and prepared by Dr. Priyanka Sahasrabudhe (Cellular Protein Chemistry, Utrecht, NL). Monoclonal mouse antibody 2.3-5 against NBD2 was kindly provided by Dr. Eric Sorscher (University of Atlanta, Emory, USA). Corrector compounds VX-809 (Vertex Pharmaceuticals or Selleck chemicals), C18 and C4 (CFFT) were dissolved in DMSO and stored at –80 °C. Proteinase K from Tritirachium album was purchased from Sigma, polyethylenimine (25 kDa branched) from Polysciences, and cycloheximide from Carl Roth.

### Expression constructs

Cloning of CFTR mutants F508del, F508del-I539T and F508del-G550E in pcDNA3.1 was described before [17]. C-terminal truncations CFTR-E395X and CFTR-E838X were constructed before [1] and subcloned from the pBS vector to pcDNA3.1 using KpnI and XhoI. CFTR and CFTR mutants G85E and F508del-CFTR in pCMVNot6.2 were a kind gift of Dr. Phil Thomas (UT Southwestern Medical Center, USA), as were CFTR mutants P67L, R74W, G85E, E92K, P205S, A455E, S549R, S549N, F508del, S1235N and S1251N in pBI CMV2. The PCR primers that were used to generate all other constructs in this study are listed in Table S1. C-terminally truncated hCFTR constructs were generated from pcDNA3.1-CFTR by PCR. PCR products were cloned into pcDNA3.1 using KpnI and XhoI. Point mutations were introduced in pcDNA3.1-CFTR by site-directed mutagenesis, after which part of the cDNA was subcloned into the pcDNA3.1-CFTR plasmid to avoid mutations in the vector backbone. All constructs were sequence verified. Construct nomenclature: N-terminal truncation ΔN35 for example starts with Methionine followed by residues 36 and following. Alternatively the construct is indicated as 36-x, with x the most C-terminal residue. C-terminal truncation D249X for example has a stop codon instead of D at position 249, and therefore has residue 248 as most C-terminal residue. In Figures 4B, C and 7B, constructs either have or lack N-terminal residues. To avoid confusion, we annotated in the Figure the N-terminal and C-terminal mutations separately: for example ΔN35-D249X then is 36-248. In the Results text we used the (incomplete) names of either the N- or the C-terminal truncations, again to avoid confusion as to which end of the construct or fragment is being described. As frequent reference is made to the Figure panels, it should be clear whether N-terminal residues are present or not.

### Transient expression and pulse-chase analysis

HEK293T cells were seeded on poly-L-Lysine (Sigma)-coated dishes to improve adherence. Twenty-four hours before the experiment, ∼40% confluent cells were transfected using the linear 25-kDa polymer polyethylenimine (1 mg/mL, pH 7.0). The DNA/PEI mixtures (ratio 1:3 (w/w), 12.5 μg PEI for a 6 cm dish) were pre-incubated for 20 minutes at RT in 150 mM NaCl. Transfected HeLa or HEK293T cells at 70-80% confluency were used for pulse-chase analysis performed essentially as described before [34]. In brief, HeLa cells were starved in MEM (ICN Biomedicals) or HEK293T cells in DMEM (Invitrogen) without cysteine and methionine for 20 minutes. Cells were pulse-labeled for the indicated times with 143 µCi/dish Easytag^TM 35^S Express Protein Labeling Mix (Perkin Elmer). After the pulse, cells were chased for indicated times in complete medium supplemented with 5 mM unlabeled methionine and cysteine. Cells were lysed with 1% (v/v) Triton X-100 in ice-cold 20 mM MES, 100 mM NaCl, 50 mM Tris-HCl pH 7.4 (MNT) and lysates were centrifuged for 10 min at 15,000 x g at 4 °C to remove nuclei.

### In-vitro translations

DNA was transcribed using T7 RNA polymerase according to manufacturer instructions (Promega). In-vitro translations were done essentially as described using semi-intact HT1080 or HEK293T cells as a source for ER membranes [81]. In brief, the cytosol of the semi-intact cells was replaced by rabbit reticulocyte lysate (Flexi Rabbit Reticulocyte Lysate System, Promega) and mRNA was translated for 1 hour at 30 °C in presence of 10 µCi/µL Easytag^TM 35^S Express Protein Labeling Mix (Perkin Elmer). The mixtures were centrifuged for 2 min at 10,000 x g at 4 °C after inhibiting protein synthesis with 1 mM cycloheximide and translocated CFTR was retrieved from the pellet fraction, which was lysed in 1% Triton-X100 in KHM (110 mM KOAc, 20 mM HEPES, 2 mM Mg(OAc)_2_ pH 7.2).

### Limited proteolysis

The fractions of the non-denaturing detergent lysates that were subjected to limited proteolysis were incubated with different concentrations Proteinase K (Sigma), as described before [1, 17]. Limited proteolysis was stopped by 2.5 mM phenylmethylsulfonyl-fluoride (PMSF) after 15 minutes incubation on ice. The reactions were analyzed directly using 12 or 15% SDS-PAGE or were used for immunoprecipitation.

### Immunoprecipitation and SDS-PAGE

CFTR was immunoprecipitated from the non-proteolyzed non-denaturing detergent lysates using polyclonal MrPink or monoclonal 2.3-5. Protease-resistant CFTR domain-specific fragments were immunoprecipitated after limited proteolysis using ECL1, or MrPink. Hence, the (proteolyzed) lysates were transferred to Protein-A Sepharose beads (GE Healthcare) that were pre-incubated for 10 minutes with antisera at 4 °C. After 3 hours immunoprecipitation for ECL1 and overnight immunoprecipitation for MrPink the complexes were washed twice at room temperature for 20 min in the following buffers: ECL1 and 2.3-5 in 10 mM Tris-HCl pH 8.6, 300 mM NaCl, 0.05% SDS and 0.05% TX-100 and MrPink in 10 mM Tris-HCl pH 8.6, 300 mM NaCl, 0.1% SDS and 0.05% Triton X-100. The washed beads were resuspended in 10 mM Tris-HCl pH 6.8, 1 mM EDTA before sample buffer was added to a final concentration of 200 mM Tris-HCl pH 6.8, 3% SDS, 10% glycerol, 0.004% bromophenol blue, 1 mM EDTA and 25 mM DTT. Samples were heated for 5 min at 55 °C before loading on SDS-PA gel (7.5-8% for full-length CFTR, 12-15% for CFTR fragments). Gels were dried and exposed to film (Kodak Biomax MR) or using a Phosphorimager (GE Healthcare Life Sciences).

### Quantification

Lane profiles were determined using a Phosphorimager (GE Healthcare Life Sciences) and the ImageQuant software (Molecular Dynamics). Radioactive band intensities were quantified with the same software, or from unsaturated exposed films using a densitometer (Bio-Rad Laboratories) and Quantity One software (Bio-Rad Laboratories).

### Structural Analysis

Images of protein structures and models were created using UCSF Chimera [82].

## ACKNOWLEDGEMENTS

We thank Dr. Eric Sorscher (Emory University, Atlanta GA, USA) for the 2.3-5 antibody, Dr. Priyanka Sahasrabudhe (Cellular Protein Chemistry, Utrecht University) for designing and preparing the E1-22 antibody, the Cystic Fibrosis Foundation (CFF) for C4 and C18, Dr. Neil Bulleid (University of Glasgow, UK) for the HT1080 fibrosarcoma cells, and all members of the Braakman-Van der Sluijs lab for effective discussions. We thank Dr. Joseline Houwman for preparing figures and editing and Dr. Hanneke Merkens-Hoelen for Figure 1A. Drs Fredrick van Goor and Jeffrey H. Stack at Vertex Pharmaceuticals are thanked for inspiring discussions and initial funding of the project. This work was funded by CFF (BRAAKM08XX0, BRAAKM14XX0), the Dutch Research Council (NWO 731.016.403, NWO 731.017.420, and NWO 022.004.019; MM), the Netherlands Organization for Health Research and Development (Zon-Mw TOP grant 40-00812-98-14103), by the Netherlands Cystic Fibrosis Foundation (NCFS, HIT-CF grant).

---

**Figure S1. NBD1 and VX-809 additively increase T1aa intensity. Related to Figure 9.**

(A) Schematic representation of the constructs used in this figure.

(B) C-terminally truncated CFTR was in-vitro translated and translocated in the presence of semi-intact cells as source of ER membranes for one hour at 30 °C in the presence of 5 μM VX-809 or vehicle (DMSO) control. Lane intensity profiles represent protease-resistant TMD1 fragments T1a (lower peak) and T1aa (upper peak) generated by proteolysis with Proteinase K and analyzed by 12% SDS-PAGE, as in Figure 9. The bar graph shows the T1aa/T1a ratio of the 100 μg/mL Proteinase-K digests.

(C) Similar to (B) and Figure 9B, but with 5 μM C18 or vehicle (DMSO) control. Protease-resistant TMD1 fragments T1a and T1aa are shown after digestion with a range of Proteinase-K concentrations and analyzed by 12% SDS-PAGE. Fragment intensity profiles are from the 100 μg/ml Proteinase-K digests.

T1a and T1aa indicate protease-resistant TMD1 fragments.

**Figure S2. Additional quantifications of T1aa/T1a ratios.**

The bar graphs show the T1aa/T1a ratio of the indicated 100 μg/mL Proteinase-K digests, except for the graph belonging to figure 8C, where the 25 μg/mL Proteinase-K digest was quantified. §, below detection level

**Supplementary Table 1.**
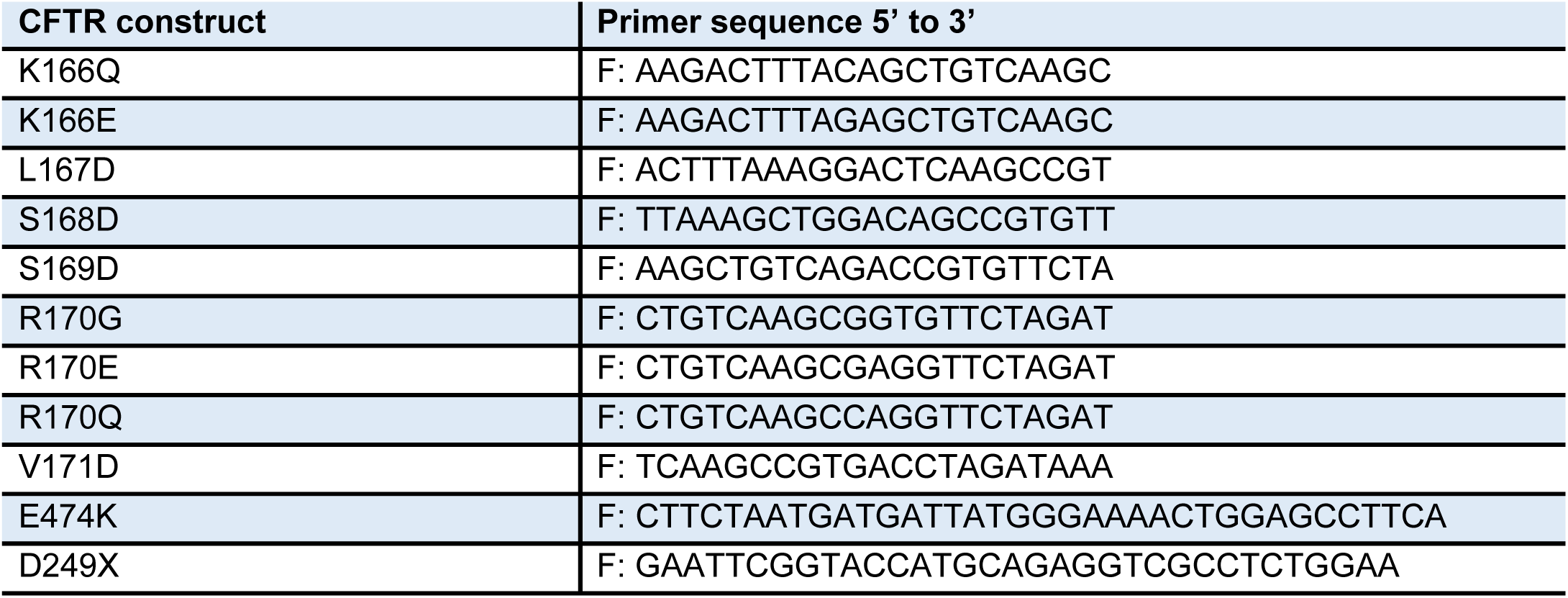
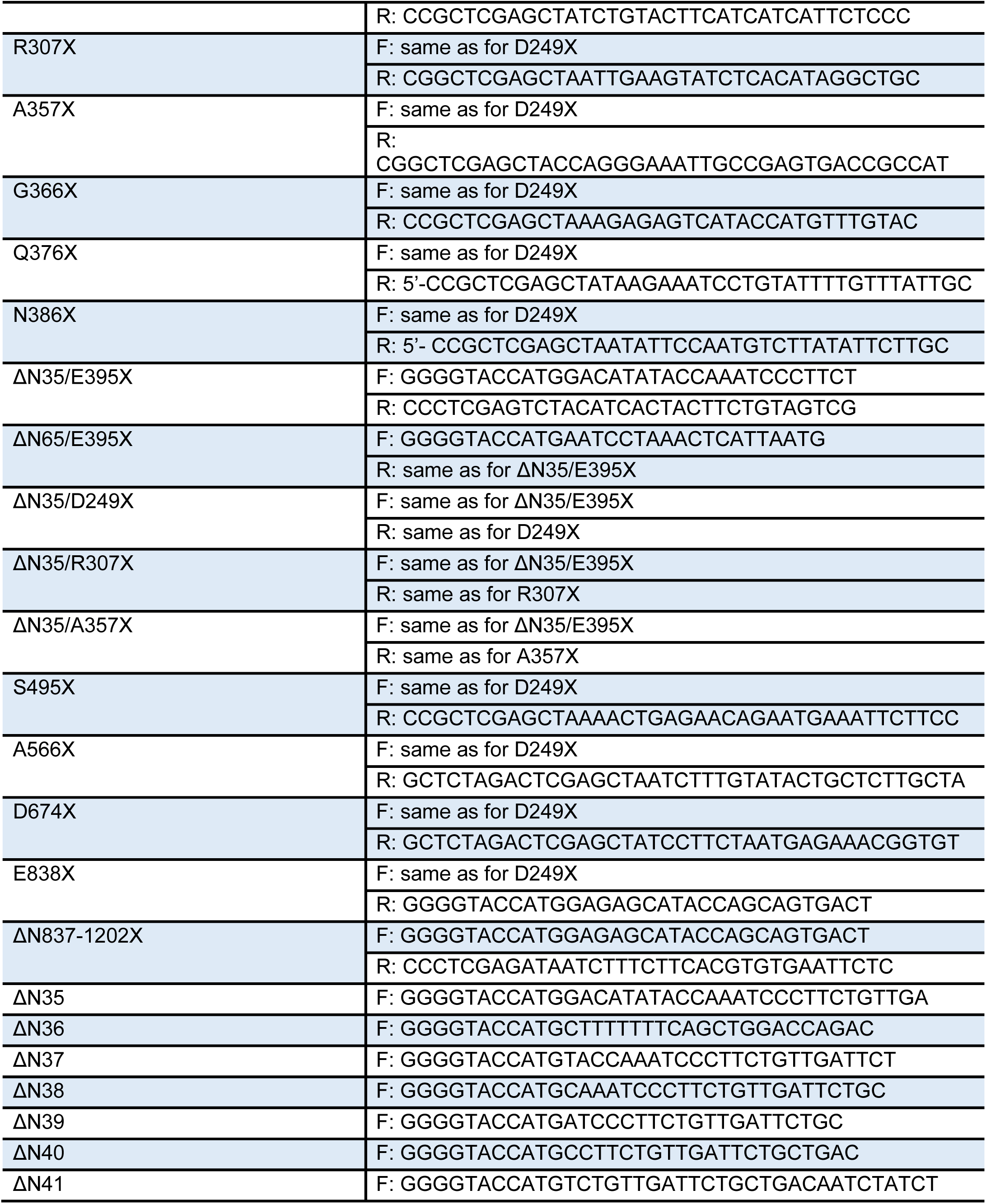
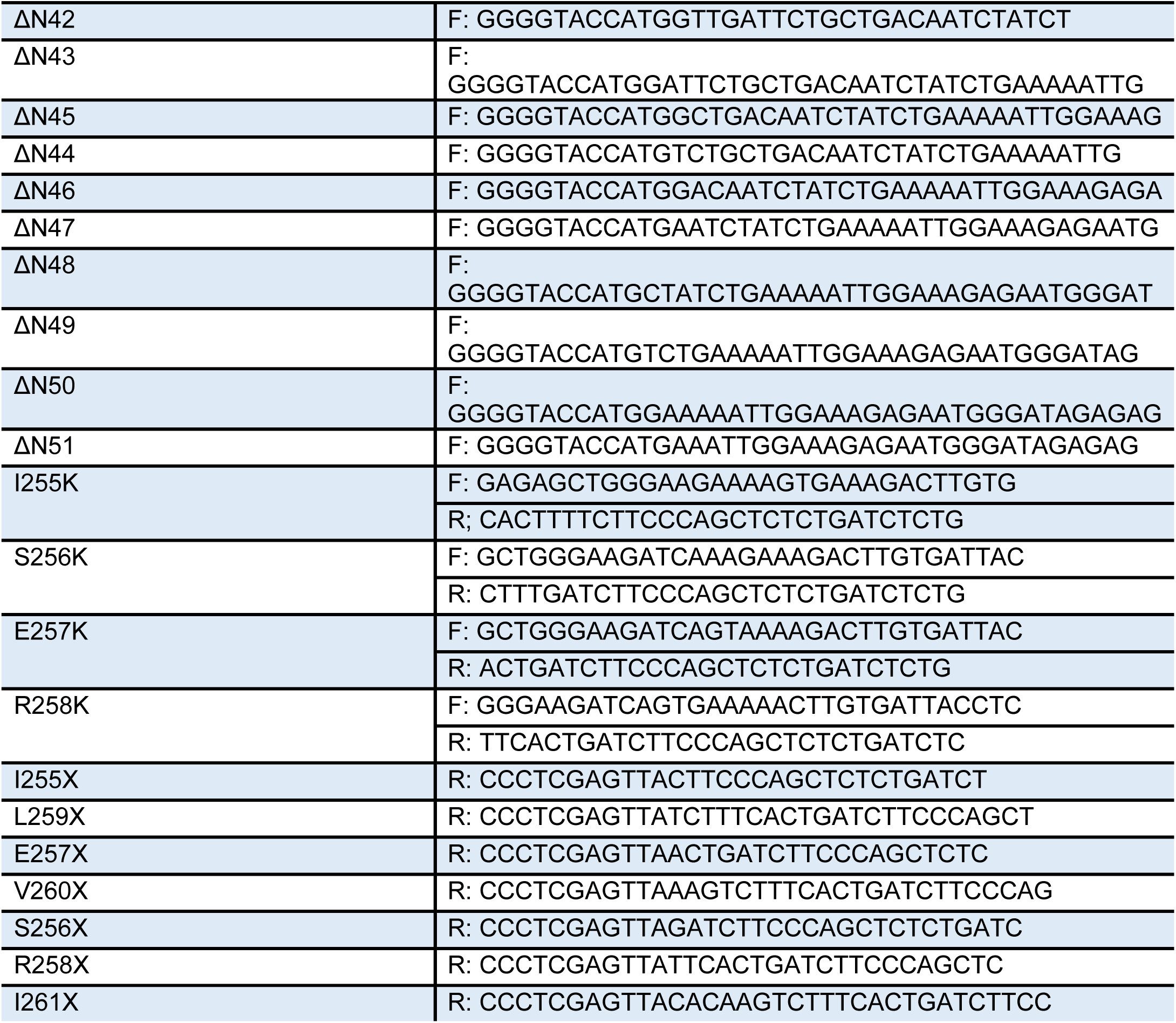
Primers used for cloning of the indicated CFTR constructs. F, forward primer; R, reverse primer.

## Notes

### Competing Interest Statement

The authors have declared no competing interest.

### Summary of Updates

Clarified a sentence; acknowledgements updated

## REFERENCES

1. Kleizen B, Van Vlijmen T, De Jonge HR, Braakman I. (2005). Folding of CFTR is predominantly cotranslational. Mol Cell. 20, 277–287.

2. Harris NJ, Reading E, Ataka K, Grzegorzewski L, Charalambous K, Liu X, et al. (2017). Structure formation during translocon-unassisted co-translational membrane protein folding. Sci Rep. 7, 8021.

3. Braakman I, Hebert DN. (2013). Protein folding in the endoplasmic reticulum. Cold Spring Harb Perspect Biol. 5, a013201.

4. Szakacs G, Abele R. (2020). An inventory of lysosomal ABC transporters. FEBS Lett.

5. Higgins CF. (1992). ABC transporters: from microorganisms to man. Annu Rev Cell Biol. 8, 67–113.

6. ter Beek J, Guskov A, Slotboom DJ. (2014). Structural diversity of ABC transporters. J Gen Physiol. 143, 419–435.

7. Anderson MP, Gregory RJ, Thompson S, Souza DW, Paul S, Mulligan RC, et al. (1991). Demonstration that CFTR is a chloride channel by alteration of its anion selectivity. Science. 253, 202–205.

8. Quinton PM, Reddy MM. (2000). CFTR, a rectifying, non-rectifying anion channel? J Korean Med Sci. 15 Suppl, S17–20.

9. Riordan JR, Rommens JM, Kerem B, Alon N, Rozmahel R, Grzelczak Z, et al. (1989). Identification of the cystic fibrosis gene: cloning and characterization of complementary DNA. Science. 245, 1066–1073.

10. Borst P, Elferink RO. (2002). Mammalian ABC transporters in health and disease. Annu Rev Biochem. 71, 537–592.

11. Kim SJ, Yoon JS, Shishido H, Yang Z, Rooney LA, Barral JM, et al. (2015). Protein folding. Translational tuning optimizes nascent protein folding in cells. Science. 348, 444–448.

12. Liu F, Zhang Z, Csanady L, Gadsby DC, Chen J. (2017). Molecular structure of the human CFTR ion channel. Cell. 169, 85–95 e88.

13. Zhang Z, Liu F, Chen J. (2017). Conformational changes of CFTR upon phosphorylation and ATP binding. Cell. 170, 483–491.

14. Qu BH, Strickland EH, Thomas PJ. (1997). Localization and suppression of a kinetic defect in cystic fibrosis transmembrane conductance regulator folding. J Biol Chem. 272, 15739–15744.

15. Qu BH, Thomas PJ. (1996). Alteration of the cystic fibrosis transmembrane conductance regulator folding pathway. J Biol Chem. 271, 7261–7264.

16. Thibodeau PH, Brautigam CA, Machius M, Thomas PJ. (2005). Side chain and backbone contributions of Phe508 to CFTR folding. Nat Struct Mol Biol. 12, 10–16.

17. Hoelen H, Kleizen B, Schmidt A, Richardson J, Charitou P, Thomas PJ, et al. (2010). The primary folding defect and rescue of ΔF508 CFTR emerge during translation of the mutant domain. PLoS One. 5, e15458.

18. Cui L, Aleksandrov L, Chang X-B, Hou Y-X, He L, Hegedus T, et al. (2007). Domain interdependence in the biosynthetic assembly of CFTR. J Mol Biol. 365, 981–994.

19. Du K, Lukacs GL. (2009). Cooperative assembly and misfolding of CFTR domains in vivo. Mol Biol Cell. 20, 1903–1915.

20. Du K, Sharma M, Lukacs GL. (2005). The ΔF508 cystic fibrosis mutation impairs domain-domain interactions and arrests post-translational folding of CFTR. Nat Struct Mol Biol. 12, 17–25.

21. Mendoza JL, Schmidt A, Li Q, Nuvaga E, Barrett T, Bridges RJ, et al. (2012). Requirements for efficient correction of ΔF508 CFTR revealed by analyses of evolved sequences. Cell. 148, 164–174.

22. Rabeh WM, Bossard F, Xu H, Okiyoneda T, Bagdany M, Mulvihill CM, et al. (2012). Correction of both NBD1 energetics and domain interface is required to restore ΔF508 CFTR folding and function. Cell. 148, 150–163.

23. Loo TW, Bartlett MC, Clarke DM. (2013). Corrector VX-809 stabilizes the first transmembrane domain of CFTR. Biochem Pharmacol. 86, 612–619.

24. Ren HY, Grove DE, De La Rosa O, Houck SA, Sopha P, Van Goor F, et al. (2013). VX-809 corrects folding defects in cystic fibrosis transmembrane conductance regulator protein through action on membrane-spanning domain 1. Mol Biol Cell. 24, 3016–3024.

25. Loo TW, Clarke DM. (2017). Corrector VX-809 promotes interactions between cytoplasmic loop one and the first nucleotide-binding domain of CFTR. Biochem Pharmacol. 136, 24–31.

26. van Willigen M, Vonk AM, Yeoh HY, Kruisselbrink E, Kleizen B, van der Ent CK, et al. (2019). Folding-function relationship of the most common cystic fibrosis-causing CFTR conductance mutants. Life Sci Alliance. 2, e201800172.

27. Cheng SH, Gregory RJ, Marshall J, Paul S, Souza DW, White GA, et al. (1990). Defective intracellular transport and processing of CFTR is the molecular basis of most cystic fibrosis. Cell. 63, 827–834.

28. Ward CL, Omura S, Kopito RR. (1995). Degradation of CFTR by the ubiquitin-proteasome pathway. Cell. 83, 121–127.

29. DeCarvalho AC, Gansheroff LJ, Teem JL. (2002). Mutations in the nucleotide binding domain 1 signature motif region rescue processing and functional defects of cystic fibrosis transmembrane conductance regulator ΔF508. J Biol Chem. 277, 35896–35905.

30. Roxo-Rosa M, Xu Z, Schmidt A, Neto M, Cai Z, Soares CM, et al. (2006). Revertant mutants G550E and 4RK rescue cystic fibrosis mutants in the first nucleotide-binding domain of CFTR by different mechanisms. Proc Natl Acad Sci U S A. 103, 17891–17896.

31. Thibodeau PH, Richardson JM, 3rd, Wang W, Millen L, Watson J, Mendoza JL, et al. (2010). The cystic fibrosis-causing mutation ΔF508 affects multiple steps in cystic fibrosis transmembrane conductance regulator biogenesis. J Biol Chem. 285, 35825–35835.

32. Van Goor F, Hadida S, Grootenhuis PD, Burton B, Stack JH, Straley KS, et al. (2011). Correction of the F508del-CFTR protein processing defect in vitro by the investigational drug VX-809. Proc Natl Acad Sci U S A. 108, 18843–18848.

33. Wainwright CE, Elborn JS, Ramsey BW. (2015). Lumacaftor-Ivacaftor in patients with cystic fibrosis homozygous for phe508del CFTR. N Engl J Med. 373, 1783–1784.

34. Braakman I, Hoover-Litty H, Wagner KR, Helenius A. (1991). Folding of influenza hemagglutinin in the endoplasmic reticulum. J Cell Biol. 114, 401–411.

35. Laselva O, Molinski S, Casavola V, Bear CE. (2018). Correctors of the major cystic fibrosis mutant interact through membrane-spanning domains. Mol Pharmacol. 93, 612–618.

36. Pedemonte N, Lukacs GL, Du K, Caci E, Zegarra-Moran O, Galietta LJ, et al. (2005). Small-molecule correctors of defective ΔF508-CFTR cellular processing identified by high-throughput screening. J Clin Invest. 115, 2564–2571.

37. Grove DE, Rosser MF, Ren HY, Naren AP, Cyr DM. (2009). Mechanisms for rescue of correctable folding defects in CFTRΔF508. Mol Biol Cell. 20, 4059–4069.

38. Okiyoneda T, Veit G, Dekkers JF, Bagdany M, Soya N, Xu H, et al. (2013). Mechanism-based corrector combination restores ΔF508-CFTR folding and function. Nat Chem Biol. 9, 444–454.

39. Peters KW, Okiyoneda T, Balch WE, Braakman I, Brodsky JL, Guggino WB, et al. (2011). CFTR Folding Consortium: methods available for studies of CFTR folding and correction. Methods Mol Biol. 742, 335–353.

40. Whittaker RG, Manthey MK, Le Brocque DS, Hayes PJ. (1994). A microtiter plate assay for the characterization of serine proteases by their esterase activity. Anal Biochem. 220, 238–243.

41. Zhang Z, Chen J. (2016). Atomic structure of the cystic fibrosis transmembrane conductance regulator. Cell. 167, 1586–1597.e1589.

42. Mornon J-P, Lehn P, Callebaut I. (2009). Molecular models of the open and closed states of the whole human CFTR protein. Cell Mol Life Sci. 66, 3469–3486.

43. Hoffmann B, Elbahnsi A, Lehn P, Decout JL, Pietrucci F, Mornon JP, et al. (2018). Combining theoretical and experimental data to decipher CFTR 3D structures and functions. Cell Mol Life Sci. 75, 3829–3855.

44. Sosnay PR, Siklosi KR, Van Goor F, Kaniecki K, Yu H, Sharma N, et al. (2013). Defining the disease liability of variants in the cystic fibrosis transmembrane conductance regulator gene. Nat Genet. 45, 1160–1167.

45. Xiong X, Bragin A, Widdicombe JH, Cohn J, Skach WR. (1997). Structural cues involved in endoplasmic reticulum degradation of G85E and G91R mutant cystic fibrosis transmembrane conductance regulator. J Clin Invest. 100, 1079–1088.

46. Patrick AE, Karamyshev AL, Millen L, Thomas PJ. (2011). Alteration of CFTR transmembrane span integration by disease-causing mutations. Mol Biol Cell. 22, 4461–4471.

47. Frydman J, Erdjument-Bromage H, Tempst P, Hartl FU. (1999). Co-translational domain folding as the structural basis for the rapid de novo folding of firefly luciferase. Nat Struct Biol. 6, 697–705.

48. Ward CL, Kopito RR. (1994). Intracellular turnover of cystic fibrosis transmembrane conductance regulator. Inefficient processing and rapid degradation of wild-type and mutant proteins. J Biol Chem. 269, 25710–25718.

49. Feng X, Barth P. (2016). A topological and conformational stability alphabet for multipass membrane proteins. Nat Chem Biol. 12, 167–173.

50. Naren AP, Cormet-Boyaka E, Fu J, Villain M, Blalock JE, Quick MW, et al. (1999). CFTR chloride channel regulation by an interdomain interaction. Science. 286, 544–548.

51. Cymer F, von Heijne G. (2013). Cotranslational folding of membrane proteins probed by arrest-peptide-mediated force measurements. Proc Natl Acad Sci U S A. 110, 14640–14645.

52. Fay JF, Aleksandrov LA, Jensen TJ, Cui LL, Kousouros JN, He L, et al. (2018). Cryo-EM visualization of an active high open probability CFTR anion channel. Biochemistry. 57, 6234–6246.

53. Zhang Z, Liu F, Chen J. (2018). Molecular structure of the ATP-bound, phosphorylated human CFTR. Proc Natl Acad Sci U S A. 115, 12757–12762.

54. Krishna MM, Englander SW. (2005). The N-terminal to C-terminal motif in protein folding and function. Proc Natl Acad Sci U S A. 102, 1053–1058.

55. Linsdell P, Tabcharani JA, Rommens JM, Hou YX, Chang XB, Tsui LC, et al. (1997). Permeability of wild-type and mutant cystic fibrosis transmembrane conductance regulator chloride channels to polyatomic anions. J Gen Physiol. 110, 355–364.

56. Tector M, Hartl FU. (1999). An unstable transmembrane segment in the cystic fibrosis transmembrane conductance regulator. EMBO J. 18, 6290–6298.

57. Wang W, Roessler BC, Kirk KL. (2014). An electrostatic interaction at the tetrahelix bundle promotes phosphorylation-dependent cystic fibrosis transmembrane conductance regulator (CFTR) channel opening. J Biol Chem. 289, 30364–30378.

58. Patrick AE, Thomas PJ. (2012). Development of CFTR Structure. Front Pharmacol. 3, 162.

59. Dawson RJ, Locher KP. (2006). Structure of a bacterial multidrug ABC transporter. Nature. 443, 180–185.

60. Locher KP. (2016). Mechanistic diversity in ATP-binding cassette (ABC) transporters. Nat Struct Mol Biol. 23, 487–493.

61. Khushoo A, Yang Z, Johnson AE, Skach WR. (2011). Ligand-driven vectorial folding of ribosome-bound human CFTR NBD1. Mol Cell. 41, 682–692.

62. He L, Aleksandrov AA, Serohijos AW, Hegedus T, Aleksandrov LA, Cui L, et al. (2008). Multiple membrane-cytoplasmic domain contacts in the cystic fibrosis transmembrane conductance regulator (CFTR) mediate regulation of channel gating. J Biol Chem. 283, 26383–26390.

63. Ehrhardt A, Chung WJ, Pyle LC, Wang W, Nowotarski K, Mulvihill CM, et al. (2016). Channel Gating Regulation by the Cystic Fibrosis Transmembrane Conductance Regulator (CFTR) First Cytosolic Loop. J Biol Chem. 291, 1854–1865.

64. Serohijos AW, Hegedus T, Aleksandrov AA, He L, Cui L, Dokholyan NV, et al. (2008). Phenylalanine-508 mediates a cytoplasmic-membrane domain contact in the CFTR 3D structure crucial to assembly and channel function. Proc Natl Acad Sci U S A. 105, 3256–3261.

65. Kanelis V, Hudson RP, Thibodeau PH, Thomas PJ, Forman-Kay JD. (2010). NMR evidence for differential phosphorylation-dependent interactions in WT and ΔF508 CFTR. EMBO J. 29, 263–277.

66. Sabusap CM, Wang W, McNicholas CM, Chung WJ, Fu L, Wen H, et al. (2016). Analysis of cystic fibrosis-associated P67L CFTR illustrates barriers to personalized therapeutics for orphan diseases. JCI Insight. 1, e86581.

67. Sharma H, Jollivet Souchet M, Callebaut I, Prasad R, Becq F. (2015). Function, pharmacological correction and maturation of new Indian CFTR gene mutations. J Cyst Fibros. 14, 34–41.

68. Veit G, Avramescu RG, Chiang AN, Houck SA, Cai Z, Peters KW, et al. (2016). From CFTR biology toward combinatorial pharmacotherapy: expanded classification of cystic fibrosis mutations. Mol Biol Cell. 27, 424–433.

69. Bartoszewski RA, Jablonsky M, Bartoszewska S, Stevenson L, Dai Q, Kappes J, et al. (2010). A synonymous single nucleotide polymorphism in ΔF508 CFTR alters the secondary structure of the mRNA and the expression of the mutant protein. J Biol Chem. 285, 28741–28748.

70. Lazrak A, Fu L, Bali V, Bartoszewski R, Rab A, Havasi V, et al. (2013). The silent codon change I507-ATC->ATT contributes to the severity of the ΔF508 CFTR channel dysfunction. FASEB J. 27, 4630–4645.

71. Lukacs GL, Chang XB, Bear C, Kartner N, Mohamed A, Riordan JR, et al. (1993). The ΔF508 mutation decreases the stability of cystic fibrosis transmembrane conductance regulator in the plasma membrane. Determination of functional half-lives on transfected cells. J Biol Chem. 268, 21592–21598.

72. Lukacs GL, Chang XB, Kartner N, Rotstein OD, Riordan JR, Grinstein S. (1992). The cystic fibrosis transmembrane regulator is present and functional in endosomes. Role as a determinant of endosomal pH. J Biol Chem. 267, 14568–14572.

73. Mijnders M, Kleizen B, Braakman I. (2017). Correcting CFTR folding defects by small-molecule correctors to cure cystic fibrosis. Curr Opin Pharmacol. 34, 83–90.

74. Gulyas-Kovacs A. (2012). Integrated analysis of residue coevolution and protein structure in ABC transporters. PLoS One. 7, e36546.

75. He L, Kota P, Aleksandrov AA, Cui L, Jensen T, Dokholyan NV, et al. (2013). Correctors of ΔF508 CFTR restore global conformational maturation without thermally stabilizing the mutant protein. FASEB J. 27, 536–545.

76. Farinha CM, King-Underwood J, Sousa M, Correia AR, Henriques BJ, Roxo-Rosa M, et al. (2013). Revertants, low temperature, and correctors reveal the mechanism of F508del-CFTR rescue by VX-809 and suggest multiple agents for full correction. Chem Biol. 20, 943–955.

77. Molinski SV, Shahani VM, Subramanian AS, MacKinnon SS, Woollard G, Laforet M, et al. (2018). Comprehensive mapping of cystic fibrosis mutations to CFTR protein identifies mutation clusters and molecular docking predicts corrector binding site. Proteins. 86, 833–843.

78. Kleizen B, Hunt JF, Callebaut I, Hwang TC, Sermet-Gaudelus I, Hafkemeyer S, et al. (2020). CFTR: New insights into structure and function and implications for modulation by small molecules. J Cyst Fibros. 19 Suppl 1, S19–S24.

79. Eckford PD, Li C, Ramjeesingh M, Bear CE. (2012). Cystic fibrosis transmembrane conductance regulator (CFTR) potentiator VX-770 (ivacaftor) opens the defective channel gate of mutant CFTR in a phosphorylation-dependent but ATP-independent manner. J Biol Chem. 287, 36639–36649.

80. Bagdany M, Veit G, Fukuda R, Avramescu RG, Okiyoneda T, Baaklini I, et al. (2017). Chaperones rescue the energetic landscape of mutant CFTR at single molecule and in cell. Nat Commun. 8, 398.

81. Wilson R, Allen AJ, Oliver J, Brookman JL, High S, Bulleid NJ. (1995). The translocation, folding, assembly and redox-dependent degradation of secretory and membrane proteins in semi-permeabilized mammalian cells. Biochem J. 307 (Pt 3), 679–687.

82. Pettersen EF, Goddard TD, Huang CC, Couch GS, Greenblatt DM, Meng EC, et al. (2004). UCSF Chimera--a visualization system for exploratory research and analysis. J Comput Chem. 25, 1605–1612.

83. Szewczyk P, Tao H, McGrath AP, Villaluz M, Rees SD, Lee SC, et al. (2015). Snapshots of ligand entry, malleable binding and induced helical movement in P-glycoprotein. Acta Crystallographica Section D. 71, 732–741.

84. Hohl M, Briand C, Grütter MG, Seeger MA. (2017). Crystal structure of a heterodimeric ABC transporter in its inward-facing conformation. Nature structural & molecular biology. 19, 395–402.

